# Quantifying the shape of cells - from Minkowski tensors to p-atic orders

**DOI:** 10.1101/2025.01.03.631196

**Authors:** Lea Happel, Griseldis Oberschelp, Valeriia Grudtsyna, Harish P. Jain, Rastko Sknepnek, Amin Doostmohammadi, Axel Voigt

## Abstract

*P*-atic liquid crystal theories offer new perspectives on how cells self-organize and respond to mechanical cues. Understanding and quantifying the underlying orientational orders is therefore essential for unraveling the physical mechanisms that govern tissue dynamics. Due to the deformability of cells this requires quantifying their shape. We introduce rigorous mathematical tools and a reliable framework for such shape analysis. Applying this to segmented cells in MDCK monolayers and computational approaches for active vertex models and multiphase field models allows to demonstrate independence of shape measures and the presence of various *p*-atic orders at the same time. This challenges previous findings and opens new pathways for understanding the role of orientational symmetries and *p*-atic liquid crystal theories in tissue mechanics and development.

## Introduction

The importance of orientational order in biological systems is becoming increasingly clear, as it plays a critical role in processes such as tissue morphogenesis, collective cell motion, and cellular extrusion. Orientational order results from the shapes of cells and their alignments with neighboring cells. Disruptions in this order, known as topological defects, are often linked to key biological events. For instance, defects—points or lines where the order breaks down—can drive cell extrusion (***Saw et al., 2017***; ***Monfared et al., 2023***) or trigger morphological changes in tissues (***Maroudas-Sacks et al., 2021***; ***Ravichandran et al., 2024***). Orientational order is linked to liquid crystal theories (***de Gennes and Prost, 1993***) and should here be interpreted in a broad sense. Recent evidence extends beyond nematic order, characterized by symmetry under 180^°^= 2*π*/2 rotation, to higher-order symmetries such as tetratic order (90^°^= 2*π*/4), hexatic order (60^°^= 2*π*/6), and even general *p*-atic orders (2*π*/*p, p* being an integer). In biological contexts, nematic order (*p* = 2) has been widely studied in epithelial tissues (***Duclos et al., 2017***; ***Saw et al., 2017***; ***Kawaguchi et al., 2017***), linking defects to cellular behaviors and tissue organization. More complex orders, such as tetratic order (*p* = 4) (***Cislo et al., 2023***) and hexatic oder (*p* = 6) (***Li and Ciamarra, 2018***; ***Durand and Heu, 2019***; ***Pasupalak et al., 2020***; ***Armengol-Collado et al., 2023***; ***Eckert et al., 2023***), have also been observed in experimental systems, offering new perspectives on how cells self-organize and respond to mechanical cues. Understanding and quantifying orientational order and the corresponding liquid crystal theory are therefore essential for unraveling the physical mechanisms that govern biological dynamics.

In physics, *p*-atic liquid crystals illustrate how particle shape and symmetry influence phase behavior, with seminal works dating back to Onsager’s theories (***Onsager, 1949***). Most prominently, hexatic order (*p* = 6) has been postulated and found by experiments and simulations as an intermediate state between crystalline solid and isotropic liquid in (***Halperin and Nelson, 1978***; ***Nelson and Halperin, 1979***; ***Murray and Van Winkle, 1987***; ***Bladon and Frenkel, 1995***; ***Zahn et al., 1999***; ***Gasser et al., 2010***; ***Bernard and Krauth, 2011***). Other examples are colloidal systems for triadic platelets (***Bowick et al., 2017***) or cubes (***Wojciechowski and Frenkel, 2004***), which lead to *p*-atic order with *p* = 3 and *p* = 4, respectively. Even pentatic (*p* = 5) and heptatic (*p* = 7) liquid crystals have been engineered (***Wang and Mason, 2018***; ***Yu and Mason, 2023***). The corresponding liquid crystal theories for *p*-atic order have only recently been defined (***Giomi et al., 2022a***; ***Krommydas et al., 2023***) and can also be used in biological contexts. However, while in colloidal systems particle shapes remain fixed, in biological tissues, cells are dynamic: their shapes are irregular, variable, and influenced by internal and external forces. These unique properties make quantifying *p*-atic order in tissues significantly more challenging, as quantification of the cell shapes is required. We will demonstrate that existing methods, such as bond-orientational order (***Nelson and Halperin, 1979***) or polygonal shape analysis (***Armengol-Collado et al., 2023***), might fail to capture the nuances of irregular cell shapes, which has severe consequences on the definition of *p*-atic order.

To address this, we adapt Minkowski tensors (***Mecke, 2000***; ***Schröder-Turk et al., 2013***)—rigorous mathematical tools for shape analysis—to quantify *p*-atic orders in cell monolayers. Minkowski tensors provide robust and sensitive measures of shape anisotropy and orientation, accommodating both smooth and polygonal shapes while remaining resilient to small perturbations. By applying these tools, we re-examine previous conclusions about *p*-atic orders in epithelial tissues and demonstrate that certain widely accepted results require reconsideration.

Our study leverages a combination of experimental and computational approaches. Experimentally, we analyze confluent monolayers of MDCK (Madin-Darby Canine Kidney) cells. ***Figure 1*** provide a snapshot of a considered monolayer of 235 wild-type MDCK cells, together with their shape classification by Minkowski tensors. The corresponding statistical data and probability distributions of these quantities are shown in ***Figure 2***. These data indicate the presence of all *p*-atic orders at once with similar probability distributions, mean values and standard devistions. In addition to these experiments we also analyze data of MDCK cells reported in (***Armengol-Collado et al., 2023***). Computationally, we employ two complementary models for cell monolayers: the active vertex model and the multiphase field model. These approaches allow us to systematically vary parameters such as cell activity and mechanical properties, providing a comprehensive view of how *p*-atic orders emerges in different contexts.

**Figure 1.**
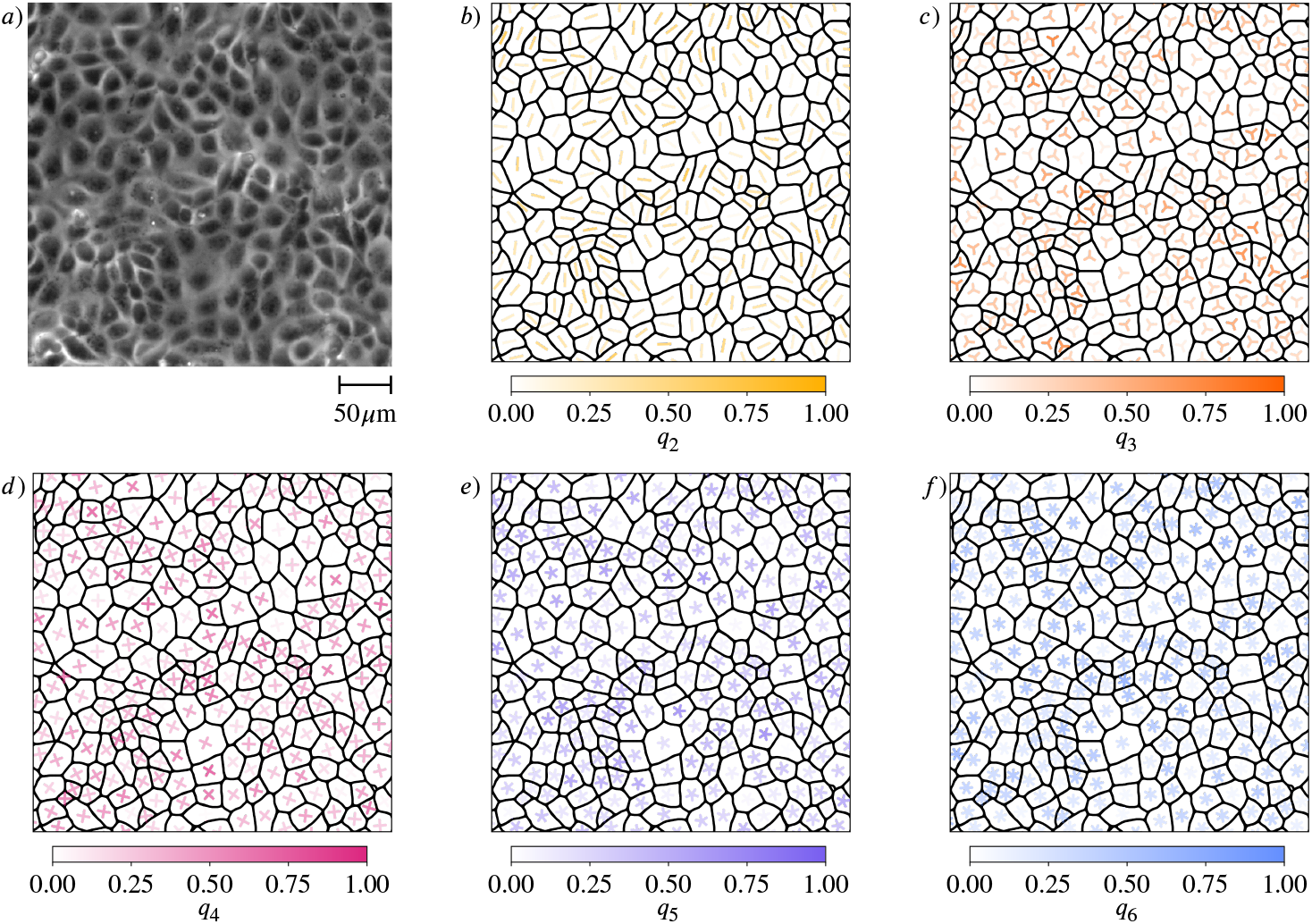
Shape classification of cells in wild-type MDCK cell monolayer. a) Raw experimental data. b) - f) Minkowski tensor, visualized using *ϑ*_*p*_ and *q*_*p*_, ***Equation 7*** (see Methods and materials) for *p* = 2, 3, 4, 5, 6, respectively. The brightness and the rotation of the *p*-atic director indicates the magnitude and the orientation, respectively. The visualization uses rotationally-symmetric direction fields (known as *p*-RoSy fields in computer graphics (***Vaxman et al., 2016***)).

**Figure 2.**
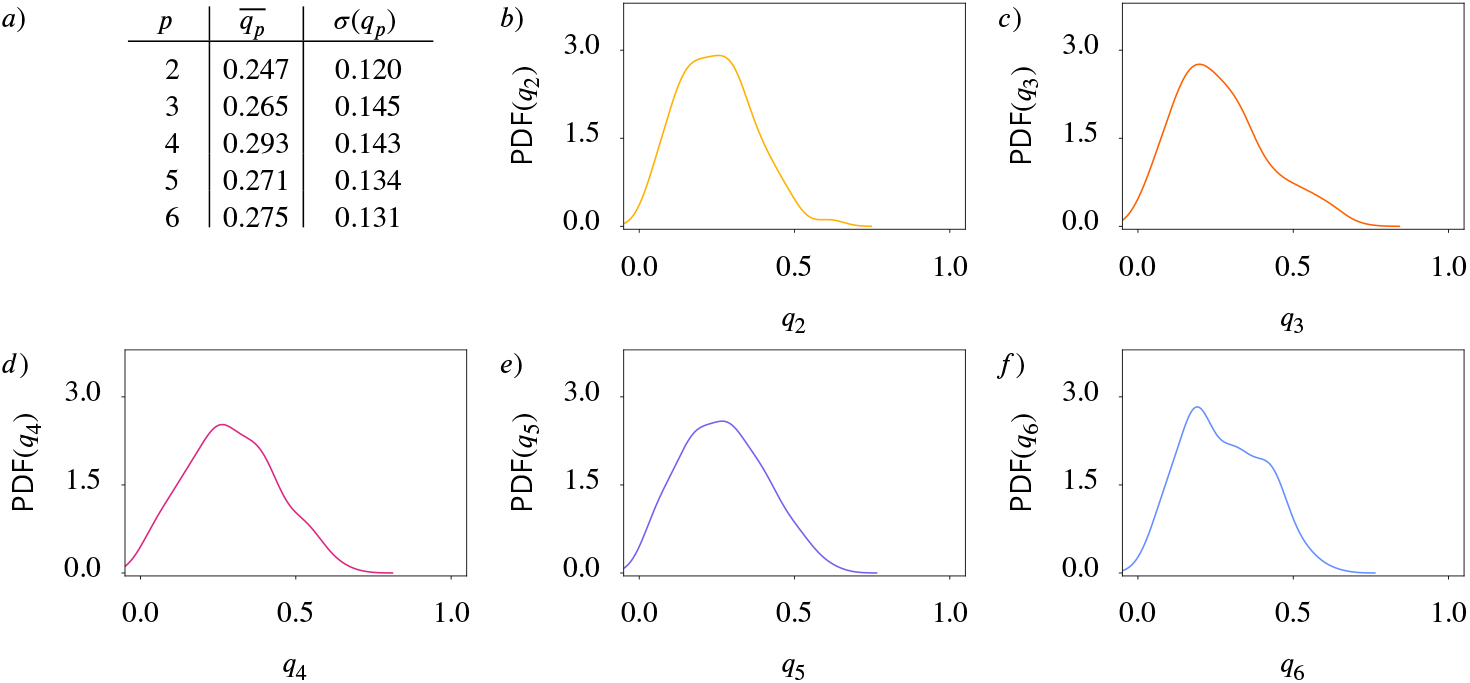
Statistical data for cell shapes identified in ***Figure 1*** (see Methods and materials). a) Mean 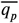 and standard deviation *σ*(*q*_*p*_) of *q*_*p*_. b) - f) Probability distribution function (PDF) of *q*_*p*_ for *p* = 2, 3, 4, 5, 6, respectively. Kde-plots are used to show the probability distribution.

By combining these robust mathematical tools, experimental insights, and computational models, we establish a reliable framework for quantifying *p*-atic orders in biological tissues. This framework not only challenges previous findings, e.g. a proposed hexatic-nematic crossover on larger length scales (***Armengol-Collado et al., 2023***), but also opens new pathways for understanding the role of orientational symmetries in tissue mechanics and development, which require to consider multiple orientational symmetries.

## Methods and materials

### Shape classification

#### Minkowski tensors

Essential properties of the geometry of a two-dimensional object are summarized by scalar-valued size measures, so called Minkowski functionals. They are defined by

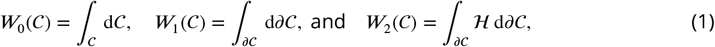

where 𝒞 is the smooth two-dimensional object, with contour *∂*𝒞 and ℋ denotes the curvature of the contour *∂*𝒞 (see ***Figure 3*** for a schematic description), as reviewed in ***Mecke (2000***). They describe the area, the perimeter and a curvature-weighted integral of the contour, respectively. Minkowski functionals are also known as intrinsic volumes. For convex objects, they have been shown to be continuous and invariant to translations and rotations. Due to these properties and Hadwiger’s characterization theorem (***Hadwiger, 1975***), they are natural size descriptors that provide essential and complete information about invariant geometric features. However, these properties also set limits to their use as shape descriptors. Due to rotation invariance, they are unable to capture the orientation of the shape. To describe more complex shape information the scalar-valued Minkowski functionals are extended to a set of tensor-valued descriptors, known as Minkowski tensors. These objects have been investigated both in mathematical (***Alesker, 1999***; ***Hug et al., 2008, 2007***; ***McMullen, 1997***) and in physical literature (***Schröder-Turk et al., 2010a***; ***Kapfer et al., 2012***). Using tensor products of the position vectors **r** and the normal vectors **n** of the contour *∂* 𝒞, defined as **r**^*a*^ ⊙ **n**^*b*^ = **r** ⊙ … ⊙ **r** ⊙ **n** ⊙ … ⊙ **n**, the first considered *a* times and the last *b* times with ⊙ denoting the symmetrized tensor product, the Minkowski tensors are defined as

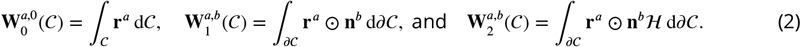

**Figure 3.**
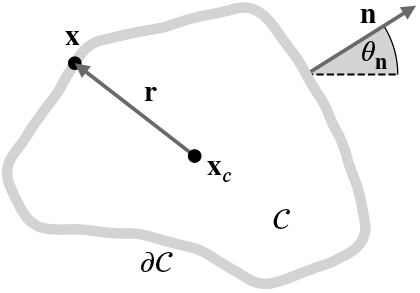
Schematic description of a two-dimensional object 𝒞 with contour *∂*𝒞. We denote the center of mass with **x**_*c*_ and vectors from **x**_*c*_ to points **x** on *∂*𝒞 with **r**. The outward-pointing normals are denoted by **n**, the corresponding angle with the *x*-axis by *θ*_**n**_.

For 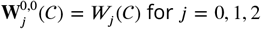, so the Minkowski tensors are extensions of the Minkowski functionals. In analogy to Minkowski functionals also Minkowski tensors have been shown to be continuous for convex objects. Also, analogies to Hadwiger’s characterization theorem exist (***Alesker, 1999***). However, for our purpose, the essential property is the continuity. It makes Minkowski functionals and Minkowski tensors a robust measure as small shape changes lead to small changes in the shape descriptor. Due to this property, Minkowski tensors have been successfully used as shape descriptors in different fields, e.g. in materials science as a robust measure of anisotropy in porous (***Schröder-Turk et al., 2010b***) and granular material (***Schröder-Turk et al., 2013***), in astrophysics to describe morphology of galaxies (***Beisbart et al., 2002***), and in biology to distinguish shapes of different types of neuronal cell networks (***Beisbart et al., 2006***) and to determine the direction of elongation in multiphase field models for epithelial tissue (***Mueller et al., 2019***; ***Wenzel and Voigt, 2021***; ***Happel and Voigt, 2024***). However, the mentioned examples exclusively use lower-rank Minkowski tensors *a* + *b* ≤ 2. Higher rank tensors have not been considered in such applications but the theory also guarantees robust description of *p*-atic orientation for *p* = 3, 4, 5, 6, …. These results hold for smooth contours but also can be extended to polyconvex shapes. Furthermore, known counter-examples for non-convex shapes have very little to no relevance in applications (***Kapfer, 2012***).

Even if all Minkowski tensors carry important geometric information, one type is particularly interesting and will be considered in the following:

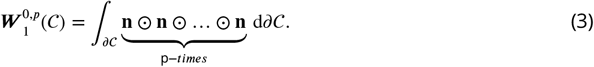

In this case, the symmetrized tensor product agrees with the classical tensor product. If applied to polygonal shapes the Minkowski tensors 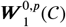 are related to the Minkowski problem for convex polytops (i.e., generalizations of three-dimensional polyhedra to an arbitrary number of dimensions), which states that convex polytops are uniquely described by the outer normals of the edges and the length of the corresponding edge (***Minkowski, 1897***). Several generalizations of this result exist (***Klain, 2004***; ***Schneider, 2013***), which makes the normal vectors a preferable quantity to describe shapes. However, Minkowski tensors contain redundant information, which asks for an irreducible representation. This can be achieved by decomposing the tensor with respect to the rotation group *SO*(2) (***Kapfer, 2012***; ***Mickel et al., 2013***; ***Klatt et al., 2022***). Following this approach, one can write

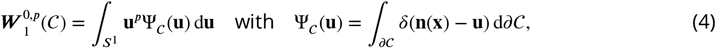

with Ψ_𝒞_ (**u**) being proportional to the probability density of the normal vectors. Identifying **u** ∈ *S*^1^ by the angle *θ* between **u** and the *x*−axis allows to write

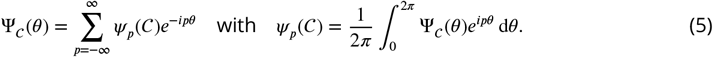

The Fourier coefficients *ψ*_*p*_(𝒞) are the irreducible representations of 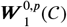 and can be written as

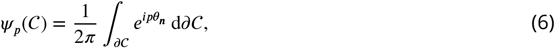

where *θ*_***n***_ is the orientation of the outward pointing normal **n**. The phase of the complex number *ψ*_*p*_(𝒞) contains information about the preferred orientation and the absolute value |*ψ*_*p*_(𝒞) | is a scalar index. We thus define

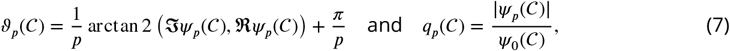

with 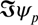 and 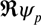 the imaginary and real part of *ψ*_*p*_, respectively, and 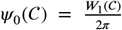. We have *q*_*p*_(𝒞) ∈ [0, 1] quantifying the strength of *p*-atic order. ***Figure 4*** illustrates the concept for polygonal and smooth shapes and ***Figure 5*** illustrates the calculation of *q*_*p*_ and *ϑ*_*p*_ at the example of an equilateral triangle. Note that the constant factor of 1/(2*π*) in ***Equation 6*** does not influence the result of ***Equation 7*** and is therefore disregarded in ***Figure 5***. Alternative derivations of ***Equation 7*** consider higher order trace-less tensors (***Virga, 2015***; ***Giomi et al., 2022a***; ***Armengol-Collado et al., 2023***) generated by the normal **n**. In this case *ϑ*_*p*_(𝒞) (orange triatic director in ***Figure 5*** right) correspond to the eigenvector to the negative eigenvalue and *ϑ*_*p*_(𝒞) − *π*/*p* (gray triatic director in ***Figure 5*** right) to the eigenvector to the positive eigenvalue. For *p* = 2 this tensor based approach corresponds to the structure tensor (***Mueller et al., 2019***).

**Figure 4.**
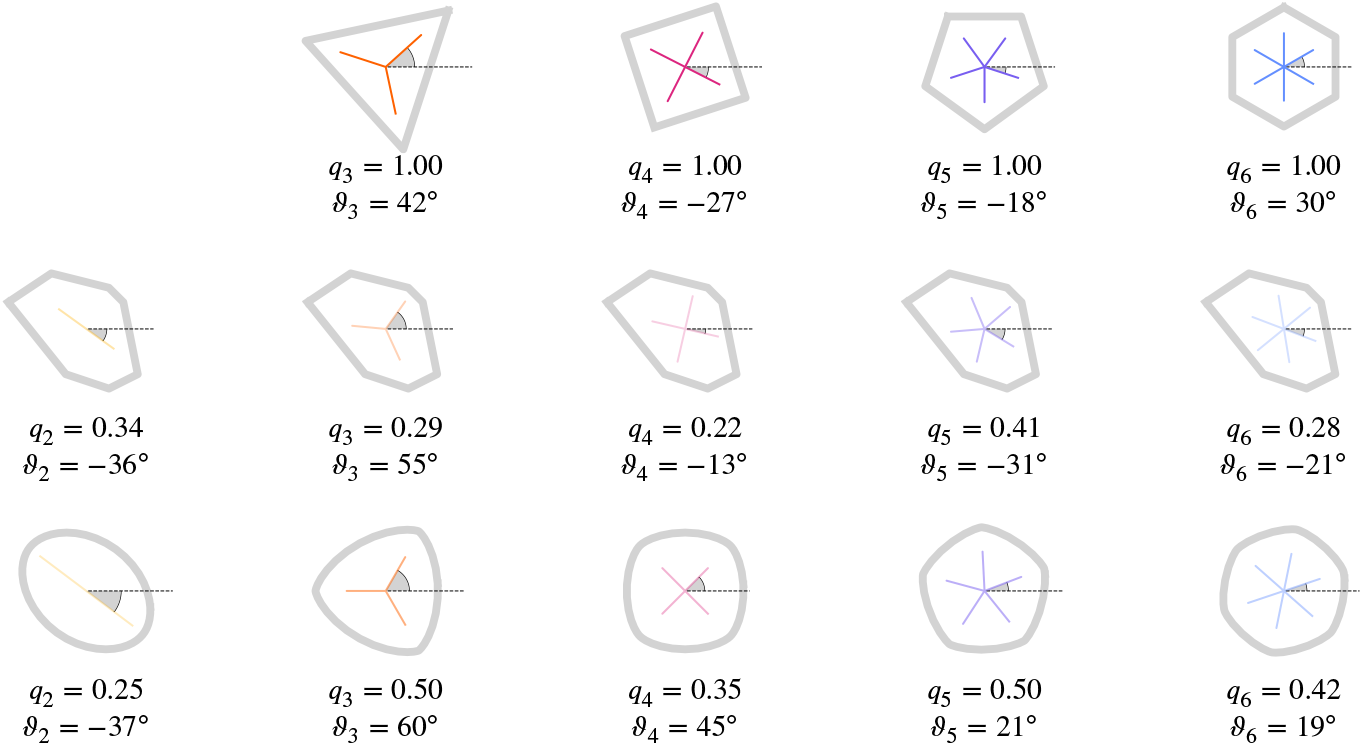
Regular and irregular shapes, adapted from ***Armengol-Collado et al. (2023***) and ***Schaller et al. (2024***), with magnitude and orientation calculated by ***Equation 7***. For regular shapes, the corresponding magnitude of *q*_*p*_ is always 1.0 and the detected angle is the minimal angle of the *p*-atic orientation with respect to the *x*-axis. Note that no shape with *q*_2_ = 1.0 is shown, as this would be a line. The visualization is according to ***Figure 1***.

**Figure 5.**
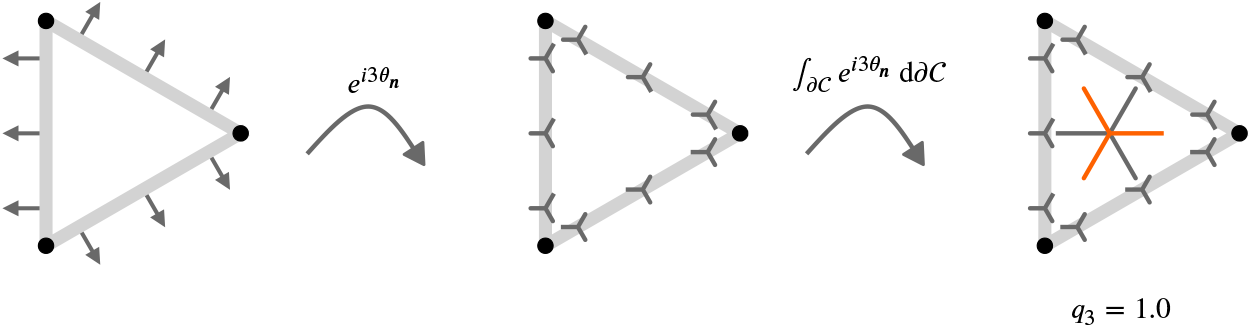
Illustrative description of the definition of *q*_3_ for an equilateral triangle. Considering rotational symmetries under a rotation 120^°^= 2*π*/3 means that vectors with an angle of 120^°^or 240^°^are treated as equal. Applied to the normals **n** (left), this means that under this rotational symmetry the normals on the three different edges are equal. Mathematically this is expressed through 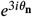 resulting in the triatic director shown instead of the normals **n** (middle). One leg of the triatic director always points in the direction as the normal. While only shown for 3 points on each edge, we obtain an orientation with the respective symmetry on every point of the contour *∂*𝒞. Considering the line integral along the contour provides the dominating triatic director, shown in the center of mass (right). To get a value between 0.0 and 1.0 for *q*_3_ we normalize this integral with the length of the contour, which corresponds to *q*_0_. As all triatic directors point in the same direction we obtain *q*_3_ = 1.0 in this specific example. To be consistent with other approaches we rotate the resulting triatic director by 60^°^= *π*/3 leading to the orange triadic director, which is the quantity used for visualization.

#### Alternative shape measures

As pointed out in the introduction, *p*-atic orders have already been identified in biological tissue and model systems. However, the way to quantify rotational symmetry in these works differs from the Minkowski tensors. In ***Armengol-Collado et al. (2023***) the cell contour is approximated by a polygon with *V* vertices having coordinates **x**_*i*_. The considered polygonal shape analysis is based on the shape function

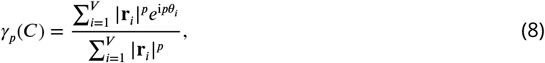

with **r**_*i*_ = **x**_*i*_ − **x**_*c*_ being the vector from the centroid **x**_*c*_ and *θ*_*i*_ being the orientation of the *i*-th vertex of the polygon with respect to its center of mass. As the centroid is computed by 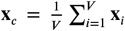 ***Armengol-Collado et al. (2023***) the only input data are the coordinates of the vertices of the *V* -sided polygon. ***Equation 8*** captures the degree of regularity of the polygon by its amplitude 0 ≤ |*γ*_*p*_| ≤ 1 and its orientation 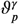 which follows as

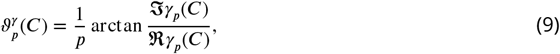

with 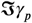 and 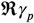 the imaginary and real part of *γ*_*p*_, respectively. Instead of the normals **n**, the descriptor is based on the position vector **r**. One might be tempted to think that ***Equation 9*** is related to the irreducible representation of 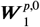. This is, however, not the case as the weighting in ***Equation 8*** is done with respect to the magnitude of **r**, while in 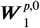 the weighting is done with respect to the contour integral. Furthermore, in contrast to the Minkowski tensors, ***Equation 8*** takes into account only the vectors at the vertices and not the vectors along the full cell contour. While this seems to be a technical detail, it has severe consequences. The theoretical basis, which guarantees continuity, no longer holds. This leads to unstable behaviour, as illustrated in ***Figure 6***. While the shape of the polygons is almost identical, there is a jump in *γ*_3_ and *γ*_4_ as we go from four to three vertices. The weighting according to the magnitude cannot cure this, as the magnitude of the shrinking vector is far from zero shortly before this vertex vanishes. In the context of cellular systems, such deformations are a regular occurrence rather than an artificially constructed test case. We will demonstrate the impact while discussing the results and recommend only using robust descriptors such as the Minkowski tensors or their irreducible representations.

**Figure 6.**
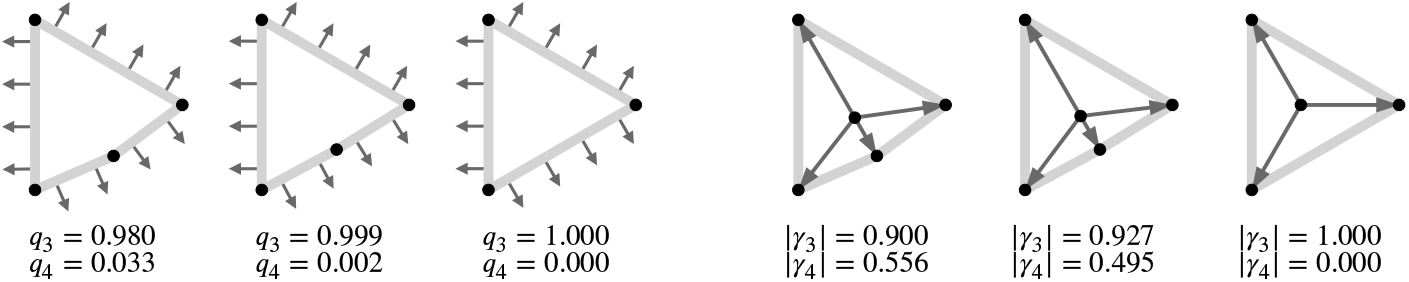
Defining *p*-atic order for deformable objects requires robust shape descriptors. Shown is the strength of *p*-atic order for a polygon converging to an equilateral triangle. a) using *q*_*p*_ and b) using *γ*_*p*_. The considered vectors used in the computations, normals **n** of the contour for the Minkowski tensors and **r**_*i*_ for *γ*_*p*_, are shown. Note that the removal of the forth vertex highly influences the value of *γ*_*p*_. How **x**_*c*_ is calculated - as the mean of the vertex coordinates or as the center of mass of the polygon - can also slightly alter the results. We here used the described approach following ***Armengol-Collado et al. (2023***).

We further note that bond order parameters 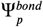, which consider the connection between cells, have also been used as shape descriptors (***Li and Ciamarra, 2018***; ***Durand and Heu, 2019***; ***Pasupalak et al., 2020***). Introduced in ***Nelson and Halperin (1979***), these parameters are

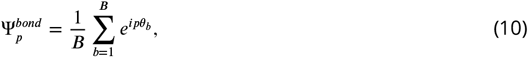

with *B* denoting the number of bonds and *θ*_*b*_ the orientation of the bond *b*. In the context of mono-layer tissues, *B* is understood as the number of neighbors, and *θ*_*b*_ as the orientation of the connection of the center of mass of the current cell with the center of mass of the neighbor *b* (***Loewe et al., 2020***; ***Monfared et al., 2023***). So cells with the same contour but different neighbor relations lead to different bond order parameters. This characteristic should already disqualify these measures as shape descriptors. However, as discussed in detail in ***Mickel et al. (2013***), they are not robust even for the task they are designed for. We therefore do not discuss them further. The same argumentation holds for other measures which are based on connectivity, as, e.g., considered in (***Graner et al., 2008***; ***Merkel et al., 2017***).

#### Coarse-grained quantities

We define coarse-grained quantities, following closely the strategy used in ***Armengol-Collado et al. (2023***). We therefore regard the coarse-grained strength of *p*-atic order *Q*_*p*_ = *Q*_*p*_(**x**), which is the average of all shape functions 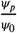 (or equivalently 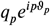) of cells whose center of mass ***x*** lies within a circle with radius *R* and center **x**. In a formula this is:

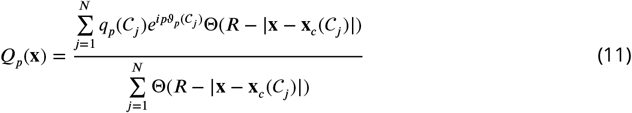

where *N* is the number of cells. 𝒞 _*i*_ denotes the *i*−th cell and Θ denotes the Heaviside step function with Θ(*x*) = 1 for *x* > 0 and Θ(*x*) = 0 otherwise. As in ***Armengol-Collado et al. (2023***) the position **x** is sampled over a square grid with a spacing close to the mean cell radius *R*_𝒞_. We calculate this as 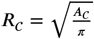 with *A*_𝒞_ the cell area. Eq. (11) provides the basis for validation of continuous *p*-atic liquid crystal theories on the tissue scale, e.g. (***Giomi et al., 2022a***,b). For comparison we also consider the coarse-grained shape function Γ_*p*_ = Γ_*p*_(**x**), which is the average of all shape functions *γ*_*p*_, which has been considered in (***Armengol-Collado et al., 2023***) and reads

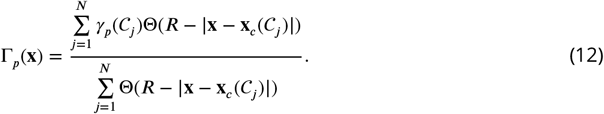

Following ***Armengol-Collado et al. (2023***) **x**_*c*_ is calculated as 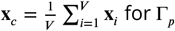.

While *Q*_*p*_ and Γ_*p*_ allow to analyze clustering on the tissue scale, their averages 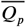 and 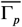 are tightly related to the statistical properties of the probability distributions of *q*_*p*_ and | *γ*_*p*_|. We consider these properties only for comparison and follow the method used in ***Armengol-Collado et al. (2023***): At first we calculate for every time instance/frame the spatial means, 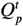 and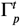, by averaging over all grid points. Then we calculate 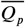 and 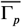 by averaging in time, so averaging over all 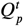 and 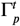. As in ***Armengol-Collado et al. (2023***) the s.e.m. and the standard derivation refer to the averaging in time. 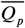 and 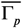 will be used for the discussion of a proposed hexatic-nematic crossover at larger length scales (***Armengol-Collado et al., 2023***).

### Simulation methods

We consider two modeling approaches, an active vertex model and a multiphase field model. Both have been proven to capture various generic properties of epithelial tissue (***Hakim and Silberzan, 2017***; ***Alert and Trepat, 2020***; ***Moure and Gomez, 2021***). The key aspects of formulations are outlined in ***Box 1*** and ***Box 2***. They refer to (***Koride et al., 2018***; ***Das et al., 2021***; ***Killeen et al., 2022***) and (***Wenzel and Voigt, 2021***; ***Jain et al., 2023, 2024b***), respectively. Parameters used within these models are listed in ***Appendix 1 Table 1*** and ***Appendix 1 Table 2***.

The initial configuration for the active vertex model simulations was built by placing *N* points randomly in a periodic simulation box of length *L* × *L* and ensuring that no two points were within a distance less than *r*_cut_ from each other. The points were used to create a Voronoi tiling. Typically, such a tiling had cells of very irregular shapes. To make cell shapes more uniform, the centroids of each tile were computed and used as seeds for a new Voronoi tiling. This process was iterated until it converged within the tolerance of 10^−5^, resulting in a so-called well-centered Voronoi tiling with cells of random shapes but similar sizes. Initial directions of polarity vectors 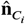 were chosen at random from a uniform distribution. The initial configuration for the multiphase field model simulations considers a regular arrangement of *N* cells of equal size in a periodic simulation box of length *L* × *L* with randomly chosen directions of self-propulsion and simulating for several time steps until the cell shapes appear sufficiently irregular. For both models we consider 100 cells and analyse time instances of the evolution.

### Experimental setup

#### Cell culture

Madin-Darby canine kidney (MDCK) cells were cultured in DMEM (DMEM, low glucose, GlutaMAX™ Supplement, pyruvate) supplemented with 10 % fetal bovine serum (FBS; Gibco) and 100 U/mL penicillin/streptomycin (Gibco) at 37 °C with 5 % CO_2_. The cell line was tested for mycoplasma.

#### Monolayer preparation

Cells were seeded on glass-bottom dishes (Mattek) pretreated with 10 µg/mL fibronectin (human plasma; Gibco) in phosphate-buffered saline (PBS, pH 7.4; Gibco). Fibronectin was incubated for 30 min at 37 °C. The initial cell seeding density was sparse. The sample was imaged approximately 24 hours later, when a confluent monolayer had formed.

#### Live cell imaging

The sample was imaged using a Nikon ECLIPSE Ti microscope equipped with an H201-K-FRAME chamber, heating system (Okolab), and CO_2_ pump (Okolab), which maintained environmental conditions at 37 °C and 5 % CO_2_. Phase-contrast images were acquired using a 10×, NA=0.3 Plan Fluor objective and an Andor Neo 5.5 sCMOS camera.

#### External dataset

As a complementary approach, we analyzed data from the study by Armengol-Collado et al. ***Armengol-Collado et al. (2023***), which included images of MDCK GII monolayers labeled for E-cadherin. These images were made publicly available by the authors via GitHub.

##### Box 1.

**Active vertex model**

**Box 1—figure 1.**
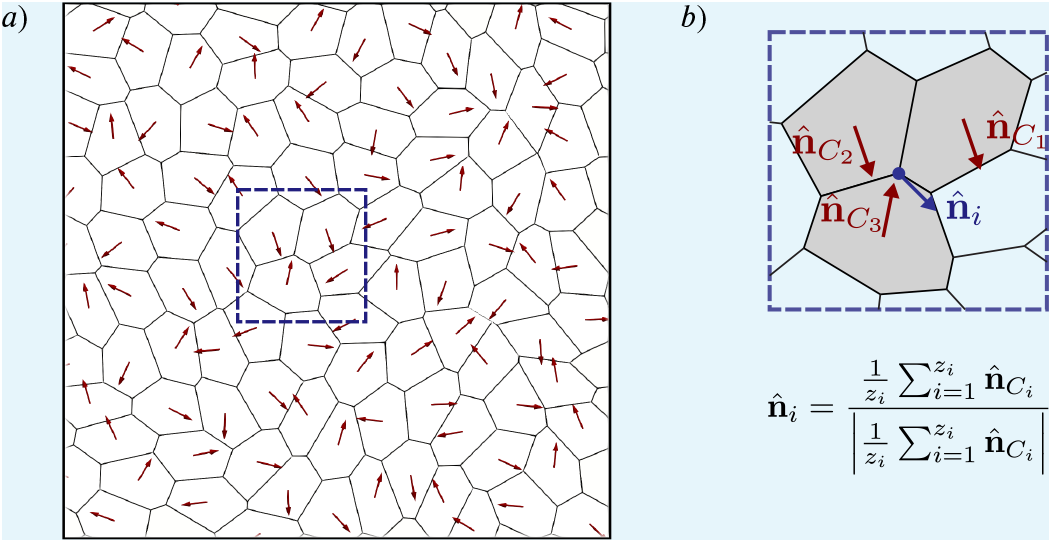
a) Cell contour of the active vertex model. Red arrows represent the polarity vectors that set each cell’s instantaneous direction of self-propulsion. b) Zoom in on a vertex surrounded by three cells showing how the direction of self-propulsion on a vertex is calculated.

The active vertex model is based on models discussed in (***Koride et al., 2018***; ***Das et al., 2021***; ***Killeen et al., 2022***). A confluent epithelial tissue is appreciated as a two-dimensional tiling of a plane with the elastic energy given as (***Honda, 1983***; ***Farhadifar et al., 2007***; ***Honda and Nagai, 2022***)

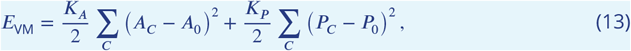

where *A*_*C*_ and *P*_*C*_ are, respectively, area and perimeter of cell *C, A*_0_ and *P*_0_ are, respectively, preferred area and perimeter, and *K*_*A*_ and *K*_*P*_ are, respectively, area and perimeter moduli. For simplicity, it is assumed that all cells have the same values of *A*_0_, *P*_0_, *K*_*A*_, and *K*_*P*_. The model can be made dimensionless by dividing ***Equation 13*** by 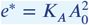,

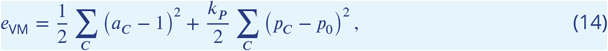

where *a*_*C*_ = *A*_*C*_ /*A*_0_ (i.e. 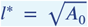 is the unit of length), 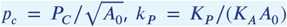, and 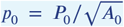 is called the shape index, and it has been shown to play a central role in determining if the tissue is in a solid or fluid state (***Bi et al., 2015***).

One can use ***Equation 13*** to find the mechanical force on a vertex *i* as 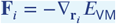, where 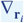 is the gradient with respect to the position **r**_*i*_ of the vertex. The expression for **F**_*i*_ is given in terms of the position of the vertex *i* and its immediate neighbours (***Tong et al., 2023***), which makes it fast to compute. Furthermore, cells move on a substrate by being noisily self-propelled along the direction of their planar polarity described by a unit-length vector 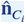 from which the polarity direction at the vertex 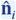 is defined, see Box 1 - figure 1 b), where *z*_*i*_ is the number of neighbours of vertex *i*. It is convenient to write 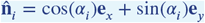, where *α*_*i*_ is an angle with the *x*−axis of the simulation box. The equations of motion for vertex *i* is a force balance between active, elastic, and frictional forces and are given as

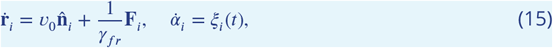

where *v*_0_ is the magnitude of the self-propulsion velocity defined as *f*_0_/*γ*_*fr*_. Furthermore the overdot denotes the time derivative, *γ*_fr_ is the friction coefficient, *f*_0_ is the magnitude of the self-propulsion force (i.e., the activity), *ξ*_*i*_(*t*) is Gaussian noise with ⟨*ξ*_*i*_(*t*)⟩ = 0 and ⟨*ξ*_*i*_(*t*)*ξ*_*j*_(*t*^′^)⟩ = 2*D*_r_*δ*_*ij*_ *δ*(*t* − *t*^′^), where *D*_r_ is the rotation diffusion constant and ⟨·⟩ is the ensemble average over the noise. Equations of motion can be non-dimensionalised by measuring time in units of *t*^*\**^ = *γ*_fr_/(*K*_*A*_*A*_0_) and force in units of 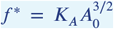 and integrated numerically using the first-order Euler-Maruyama method with timestep τ.

##### Box 2.

**Multiphase field model**

**Box 2—figure 1.**
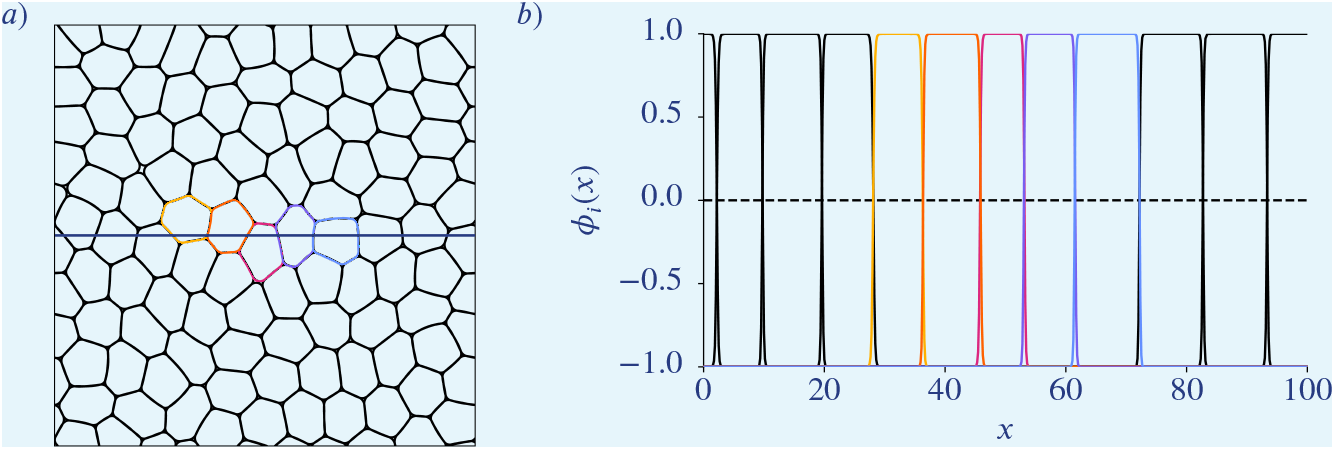
a) Cell contours of the multiphase field model. b) Corresponding phase field functions along the horizontal line in a). Colours correspond to the once in a).

The multiphase field modelling approach follows ***Wenzel and Voigt (2021***); ***Jain et al. (2023***, 2024b). Each cell is described by a scalar phase field variable *ϕ*_*i*_ with *i* = 1, 2, …, *N*, where the bulk values *ϕ*_*i*_ *≈* 1 denotes the cell interior, *ϕ*_*i*_ *≈* −1 denotes the cell exterior, and with a diffuse interface of width 𝒪 (*ϵ*) between them representing the cell boundary. The phase field *ϕ*_*i*_ follows the conservative dynamics

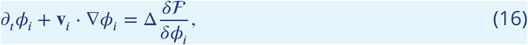

with the free energy functional ℱ = ℱ_*CH*_ + ℱ_*INT*_ containing the Cahn-Hilliard energy

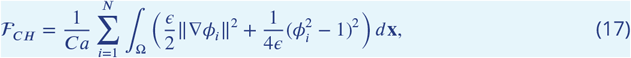

where the Capillary number *Ca* is a parameter for tuning the cell deformability, and an interaction energy consisting of repulsive and attractive parts defined as

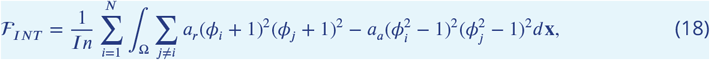

where *In* denotes the interaction strength, and *a*_*r*_ and *a*_*a*_ are parameters to tune contribution of repulsion and attraction. The repulsive term penalises overlap of cell interiors, while the attractive part promotes overlap of cell interfaces. The relation to former formulations is discussed in ***Happel and Voigt (2024***).

Each cell is self-propelled, and the cell activity is introduced through an advection term. The cell velocity field is defined as

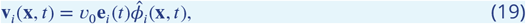

where *v*_0_ is used to tune the magnitude of activity, 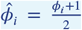 and **e**_*i*_ = [cos *θ*_*i*_(*t*), sin *θ*_*i*_(*t*)] is its direction. The migration orientation *θ*_*i*_(*t*) evolves diffusively with a drift that aligns to the principal axis of cell’s elongation as 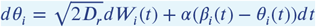, where *D*_*r*_ is the rotational diffusivity, *W*_*i*_ is the Wiener process, *β*_*i*_(*t*) is the orientation of the cell elongation and *α* controls the time scale of this alignment.

The resulting system of partial differential equations is considered on a square domain with periodic boundary conditions and is solved by the finite element method within the toolbox AMDiS (***Vey and Voigt, 2007***; ***Witkowski et al., 2015***) and the parallelization concept introduced in ***Praetorius and Voigt (2018***) is considered, which allows scaling with the number of cells. We in addition introduce a de Gennes factor in ℱ_*CH*_ to ensure *ϕ*_*i*_ ∈ [−1, 1], see ***Salvalaglio et al. (2021***).

### Image analysis

All image data, including both our phase-contrast recordings and the E-cadherin-labeled external dataset, were analyzed using Cellpose ***Stringer et al. (2021***). Segmentation was performed using manually trained models, each tailored to its respective dataset.

### Extraction of the contour

Starting out from the segmented images (stored as grayscale images) we extract the contour of the cells using the python package *scikit-image* (***van der Walt et al., 2014***). To get rid of pixel-shaped artifacts we smooth the contour by replacing every coordinate by the average over 9 neighboring points in the outline.

## Results

Quantifying orientational order in biological tissues can be realized by Minkowsky tensors. The orientation *ϑ*_*p*_ and the strength of *p*-atic order *q*_*p*_ in ***Equation 7*** can be computed for each cell. Minkowski tensors provide reliable quantities describing how cell shapes align with specific rotational symmetries. As already indicated in ***Figure 2*** situations might occur in which rotational symmetries can not be associated with one specific *p*, but various symmetries seem to be present at the same time. One might be tempted to compare the probability distribution functions (PDFs) of *q*_*p*_ or the mean values 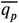 for different *p* in order to identify a dominating *p*-atic order. This is particularly important for interpreting nematic (*p* = 2) and hexatic (*p* = 6) orders, which describe distinct symmetries but have been found to coexist in biological systems. However, a direct comparison of these quantities only makes sense if they are comparable to each other. As already mentioned in ***Figure 4*** this is not the case, as, e.g., *q*_2_ = 1.0 cannot be realized for a cell with a given area. Another question one might ask is if these values are independent of each other. To answer this question, we first address statistically if *q*_2_ and *q*_6_, evaluated for each cell, are independent. Second, we explore how *q*_*p*_ depends on key parameters determining tissue mechanics. In a third step we coarsegrain these quantities using the measures *Q*_2_ and *Q*_6_ in ***Equation 11***. With these quantities we address a proposed hexatic-nematic crossover in epithelial tissue, where hexatic order dominates at small scales and nematic order prevails at larger scales (***Eckert et al., 2023***; ***Armengol-Collado et al., 2023, 2024***). For these three tasks we analyze simulation data from two computational models, the active vertex model ***Box 1*** and the multiphase field model ***Box 2***. Although these models differ conceptually, both have been validated in studies of cell monolayer mechanics, including solid-liquid transitions, neighbor exchange dynamics, and stress profiles (***Fletcher et al., 2014***; ***Alt et al., 2017***; ***Li et al., 2019***; ***Balasubramaniam et al., 2021***; ***Wenzel and Voigt, 2021***; ***Sknepnek et al., 2023***; ***Melo et al., 2023***). Key model parameters are the deformability and the activity strength. They are crucial for capturing the coarse-grained properties of confluent tissues (***Jain et al., 2023, 2024b***). In the active vertex model, deformability is controlled by the target shape index *p*_0_, reflecting the balance between cell-cell adhesion and cortical tension. The multiphase field model, by contrast, encodes deformability through the capillary number *Ca*, which directly incorporates cortical tension. We vary activity strength (*v*_0_), the shape index (*p*_0_), and the capillary number (*Ca*), ensuring all parameter combinations remain in the fluid regime. Fluidity was confirmed using by mean square displacement (MSD) (***Loewe et al., 2020***), neighbor number variance (***Wenzel and Voigt, 2021***), and the self-intermediate scattering function (***Bi et al., 2016***).

These studies demonstrate independence of *q*_2_ and *q*_6_, a general trend of increasing 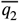 and decreasing 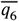 for higher activity and deformability, and a consistent decrease of 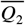 and 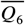 for increasing coarse-graining radius *R*. However, as *q*_2_ and *q*_6_ are not directly comparable and *q*_2_ and *q*_6_ (and therefore also *Q*_2_ and *Q*_6_) are independent, the concept of a hexatic-nematic transition - typically requiring a single order parameter - may not be applicable in this context. Even if the proposed hexatic-nematic crossover (***Armengol-Collado et al., 2023***) is not a formal phase transition, one might expect it to be quantifiable. Yet, its characterization appears to depend strongly on the maximum attainable value of *q*_2_, which in turn is influenced by several parameters. To further explore this, we reanalyze the experimental data for MDCK cells from (***Armengol-Collado et al., 2023***) using Minkowski tensors and compute *q*_2_ and *q*_6_ as defined in ***Equation 7***. Our analysis also indicates independence of *q*_2_ and *q*_6_ supporting the same interpretation as above. The statistical properties and probability distributions of these data remain consistent when full cellular boundaries from microscopy images are used. However, an increase in hexatic order (*p* = 6) is observed when cell shapes are approximated by polygons. This suggests that the dominant hexatic order reported at the cellular scale in ***Armengol-Collado et al. (2023***) may stem from the geometric simplification of cell boundaries. We also compute *Q*_2_ and *Q*_6_ (***Equation 11***) and again observed a consistent decrease with increasing coarse-graining radius *R*, without evidence of a measurable hexatic-nematic crossover. To better understand the differences with the findings in ***Armengol-Collado et al. (2023***) we further analyze both simulation and experimental data using the alternative shape measures *γ*_*p*_ (***Equation 8***) and Γ_*p*_ (***Equation 12***) considered in ***Armengol-Collado et al. (2023***). These measures reproduce the reported results.

### Independence of *q*_2_ and *q*_6_

We examine the distribution of (*q*_2_, *q*_6_) values across deformability-activity parameter pairs in both computational models (***Figure 7***). Both models show consistent trends: In near-solid regimes (***Figure 7*** lower left), *q*_2_ and *q*_6_ values cluster tightly due to restricted shape fluctuations. However, even in this regime, small *q*_2_ values can correspond to either small or large *q*_6_ values, and vice versa. In more fluid-like regimes, with higher activity and higher deformability (***Figure 7*** upper right), *q*_2_ and *q*_6_ values become highly scattered. Each *q*_2_ value spans a broad range of *q*_6_ values, and vice versa, indicating their independence. In order to quantify this we compute the distance correlation (***Székely et al., 2007***), which is a statistical measure quantifying linear and non-linear dependency in given data. Thereby a value of 0.0 corresponds to independence whereas a value of 1.0 corresponds to a strong dependence between the datasets. As can be seen in ***Figure 7—figure Supplement 1*** the obtained distance correlation for *q*_2_ and *q*_6_ is quite low, underscoring that these quantities are independent. Furthermore the corresponding P-values, as shown in ***Figure 7—figure Supplement 2*** are mostly larger than 0.1, indicating that the weak correlation found in ***Figure 7— figure Supplement 1*** is not significant. This leads to the conclusion that *q*_2_ and *q*_6_ measure distinct aspects of cell shape anisotropy. As a consequence both orders, nematic and hexatic, need to be considered independently. There cannot be a single parameter which describes a crossover between both.

**Figure 7.**
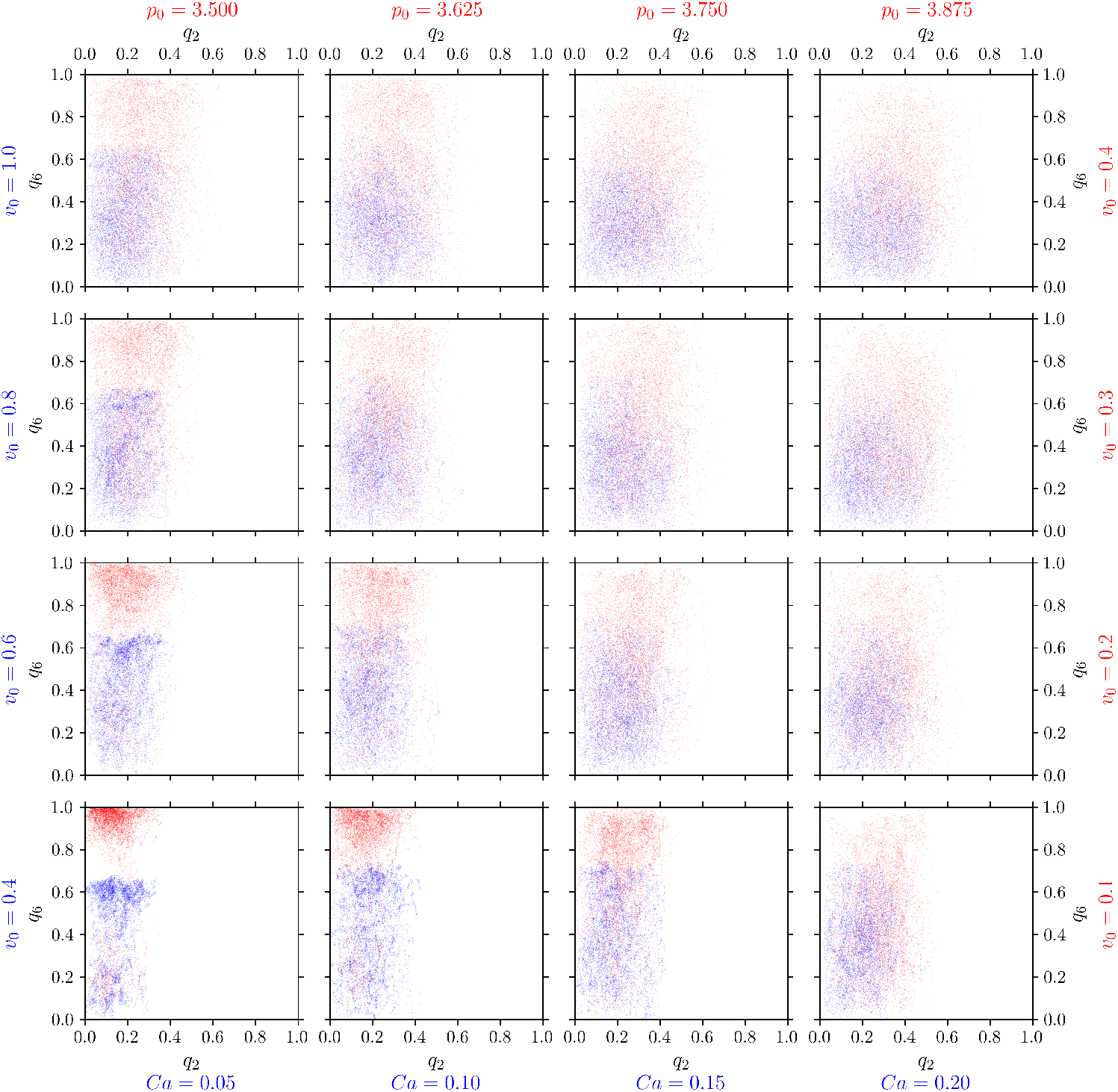
Nematic (*p* = 2) and hexatic (*p* = 6) order are independent of eachother. *q*_6_ (y-axis) versus *q*_2_ (x-axis) for all cells in the multiphase field model (blue) and active vertex model (red). For each cell and each timestep we plot one point (*q*_2_, *q*_6_). Each panel corresponds to specific model parameters; *Ca* and *v*_0_ for multiphase field model, and *p*_0_ and *v*_0_ for the active vertex model, representing deformability and activity, respectively. **Figure 7—figure supplement 1**. Distance correlation *dCor*(*q*_2_, *q*6) between *q*_2_ and *q*_6_ **Figure 7—figure supplement 2**. P-values of the distance correlation 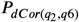 between *q*_2_ and *q*_6_

A full investigation of potential dependencies between *q*_*p*_ for arbitrary combinations of *p*’s resulting, e.g. from symmetry arguments is beyond the scope of this paper.

### Dependence of *q*_*p*_ on activity and deformability

We now explore how tissue properties such as activity and deformability influence cell shape and orientational order. We again focus on nematic (*p* = 2) and hexatic (*p* = 6) orders, as shown in ***Figure 8—figure Supplement 1*** and ***Figure 8—figure Supplement 2***. Additional results for *p* = 3, 4, 5 are provided in ***Appendix 2-Figure 14– Appendix 2-Figure 16*** (fixed activity) and ***Appendix 2-Figure 17– Appendix 2-Figure 19*** (fixed deformability). In all plots all cells and all time steps are considered. In ***Figure 8—figure Supplement 1***, we vary deformability (*p*_0_ in the active vertex model and *Ca* in the multiphase field model) while keeping the activity *v*_0_ constant. Results are presented for the active vertex model (left column) and the multiphase field model (right column). Activity increases from bottom to top rows. Both models show qualitatively similar trends in the probability distribution functions (PDFs) of *q*_2_ and *q*_6_. For *p* = 6, increasing deformability shifts the PDF of *q*_6_ to the left, indicating lower mean values. In contrast, for *p* = 2, higher deformability leads to higher *q*_2_ values, reflected in a rightward shift of the PDF. This trend is confirmed by the mean values, shown as a function of deformability (*p*_0_ and *Ca*, respectively) in the inlets. Additionally, the PDFs broaden with increasing deformability, and this effect is more pronounced at lower activity levels. A notable difference between the models is the range of *q*_*p*_ values. In the multiphase field model, *q*_*p*_ rarely exceeds 0.8 due to the smoother, more rounded cell shapes, whereas the active vertex model often produces higher *q*_*p*_ values. In ***Figure 8—figure Supplement 2***, deformability is held constant while activity *v*_0_ is varied. Results are again shown for the active vertex model (left column) and the multiphase field model (right column), with increasing activity indicated by brighter colors. Deformability increases from bottom to top rows. The trends mirror those observed in ***Figure 8— figure Supplement 1***. Increasing activity reduces the mean value of *q*_6_ while increasing *q*_2_, see inlets. The effects of activity are more pronounced at lower deformabilities; at higher deformabilities, differences between parameter regimes diminish. The overall behavior of *q*_2_ and *q*_6_ is summarized in ***Figure 8***, which shows the mean values 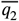 and 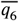 as functions of activity and deformability. For both models:

- 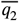 increases with higher activity or deformability
- 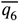 decreases with higher activity or deformability.

**Figure 8.**
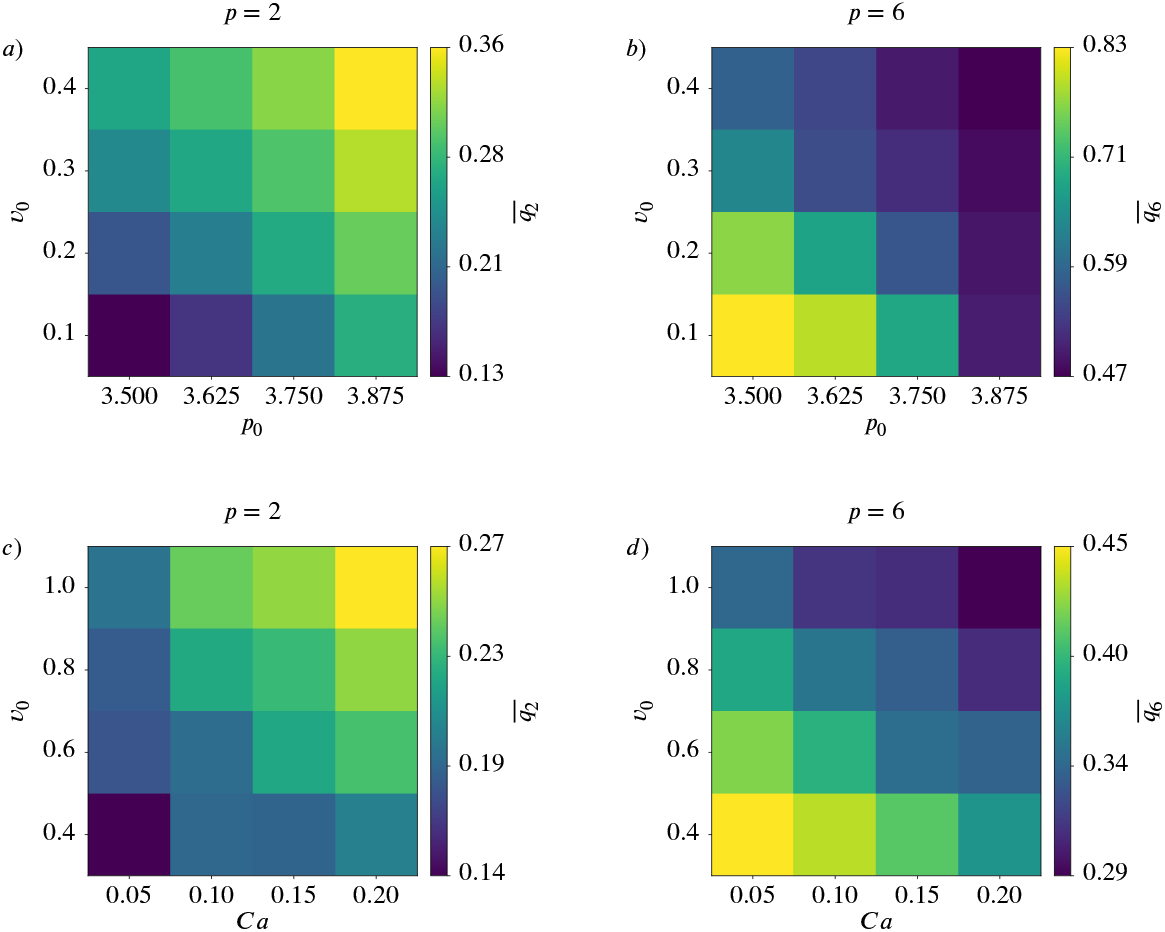
Nematic (*p* = 2) and hexatic (*p* = 6) order depend on activity and deformability of the cells. Mean value 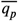 for *p* = 2 (left) and *p* = 6 (right) as function of deformability *p*_0_ or *Ca* and activity *v*_0_ for active vertex model (a) and b)) and multiphase field model (c) and d)). **Figure 8—figure supplement 1**. Nematic (*p* = 2) and hexatic (*p* = 6) order depend on deformability of the cells **Figure 8—figure supplement 2**. Nematic (*p* = 2) and hexatic (*p* = 6) order depend on activity of the cells

This behavior is consistent with a trend towards nematic order in more dynamic regimes and towards hexatic order in more constrained, less dynamic regimes. The first is further confirmed by recent findings in analysing T1 transitions and their effect on cell shapes ***Jain et al. (2024a***). These studies suggest that cells transiently elongate when they are undergoing T1 transitions. As the number of T1 transitions increases with activity or deformability ***Jain et al. (2024b***) this elongation contributes to the observed behavior. The second is consistent with the emergence of hexagonal arrangements in solid-like states.

Corresponding results for *p* = 3, 4, 5 are shown in ***Appendix 2-Figure 20***. While *q*_3_, *q*_4_, *q*_5_ also increase with increasing activity or deformability, the dependency is not as pronounced as for *q*_2_.

### Coarse-grained quantities *Q*_2_ and *Q*_6_ and potential hexatic-nematic crossover

For every parameter configuration in the active vertex model (***Figure 9—figure Supplement 1***) and in the multiphase field model (***Figure 9—figure Supplement 2***) we compute the coarse-grained quantities *Q*_2_ and *Q*_6_ for various coarse-graining radii *R*. One might ask the question if the observed trends for 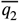 and 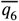 for higher activity and deformability in ***Figure 8*** are also present on larger scales. We therefore investigate the behavior of 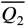 and 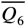 upon varying activity and deformability, see ***Figure 9***. This is exemplified for *R*/*R*_*cell*_ = 8.0. The trends seen for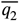 and 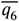 - so increasing 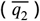 or decreasing 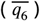 with higher activity or deformability - are lost for 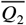 and less pronounced for 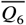.

**Figure 9.**
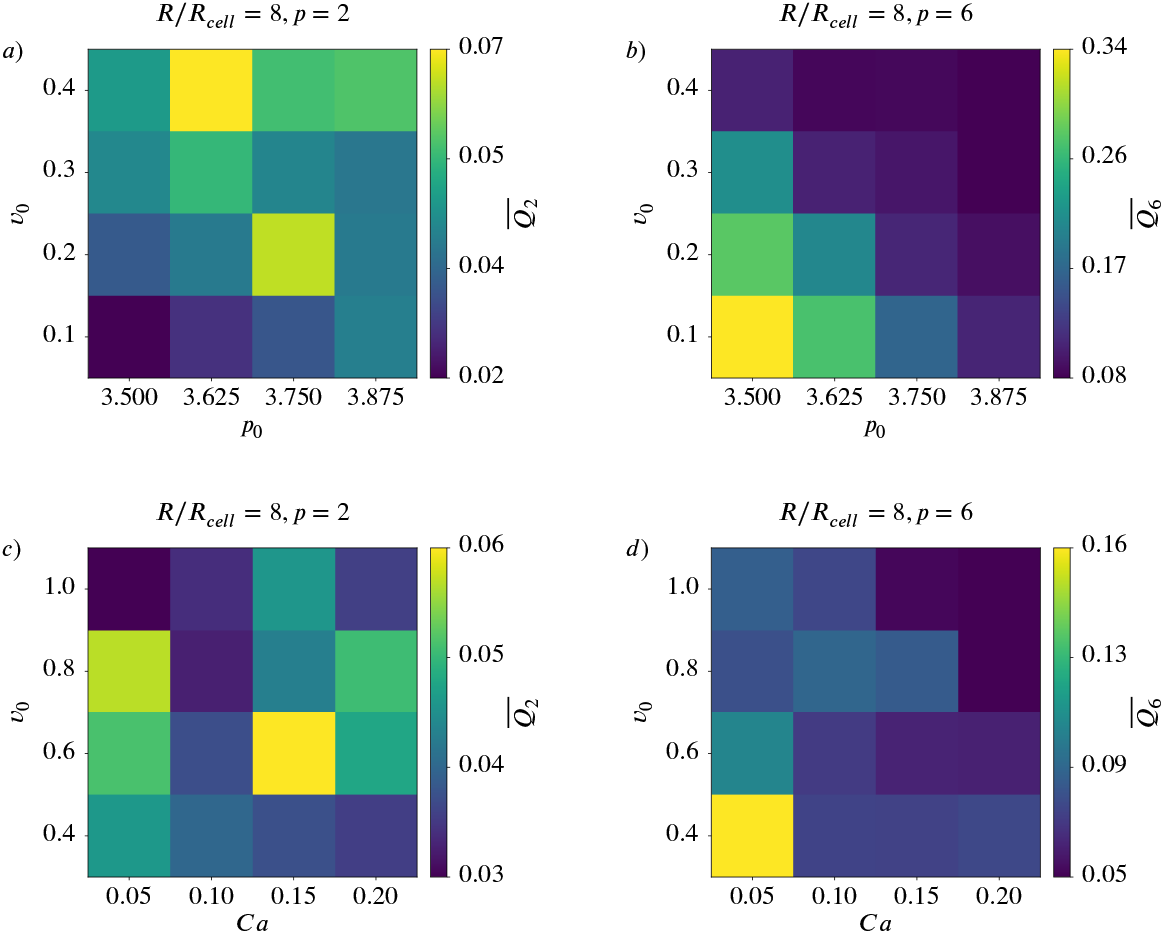
Coarse-gained nematic (*p* = 2) and hexatic (*p* = 6) order for *R*/*R*_*cell*_ = 8 depend on activity and deformability of the cells. Mean value 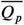 for *p* = 2 (left) and *p* = 6 (rigth) as function of deformability *p*_0_ or *Ca* and activity *v*_0_ for active vertex model (*a*) and *b*)) and multiphase field model (*c*) and *d*)). **Figure 9—figure supplement 1**. 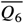 versus 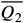 for different coarse graining radii in the active vertex model. **Figure 9—figure supplement 2**. 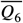 versus 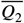 for different coarse graining radii in the multiphase field model.

In ***Armengol-Collado et al. (2023***) a similar approach was used to identify a hexatic-nematic crossover, see (***Armengol-Collado et al., 2023***, Fig. 3e). This cross-over is considered at the coarse-graining radius at which the two curves for 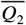 and 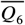 as a function of *R* cross. While already conceptually questioned above we consider these investigations to compare with ***Armengol-Collado et al. (2023***). However, regardless of the model there is no consistent trend indicative of a potential crossover. These results question the proposed hexatic-nematic crossover reported in (***Armengol-Collado et al., 2023***). To further explore this issue we next test the existence of such a crossover directly on the data considered in (***Armengol-Collado et al., 2023***).

### Analyzing experimental data for MDCK cells

The experimental data for confluent monolayers of MDCK GII cells used in (***Armengol-Collado et al., 2023***) are provided in two different formats, as microscopy images and as polygon data with the calculated vertex points per cell. We consider both formats and all 68 provided configurations. The polygonal data are directly used to compute *q*_2_ and *q*_6_. The experimental microscopy images are segmented and the extracted cell boundaries are used to compute *q*_2_ and *q*_6_.

We compute the probability distribution functions (PDFs) of *q*_2_ and *q*_6_, ***Figure 10*** a) for both data sets. While they are similar if the full cellular contour from the microscopy images is considered the PDFs strongly differ if the polygonal shapes are used. The dominating hexatic (*p* = 6) order is therefore just a consequence of the approximation of the cell boundaries by polygons. This different behavior, which shows larger values for *q*_6_ for the polygonal shapes has the same origin as the difference between the regular and rounded shapes in ***Figure 4***. Further differences result from the different accessible parameter range. While *q*_6_ = 1.0 is possible for a perfect hexagon, *q*_2_ = 1.0 cannot be realised, as this would correspond to a line.

**Figure 10.**
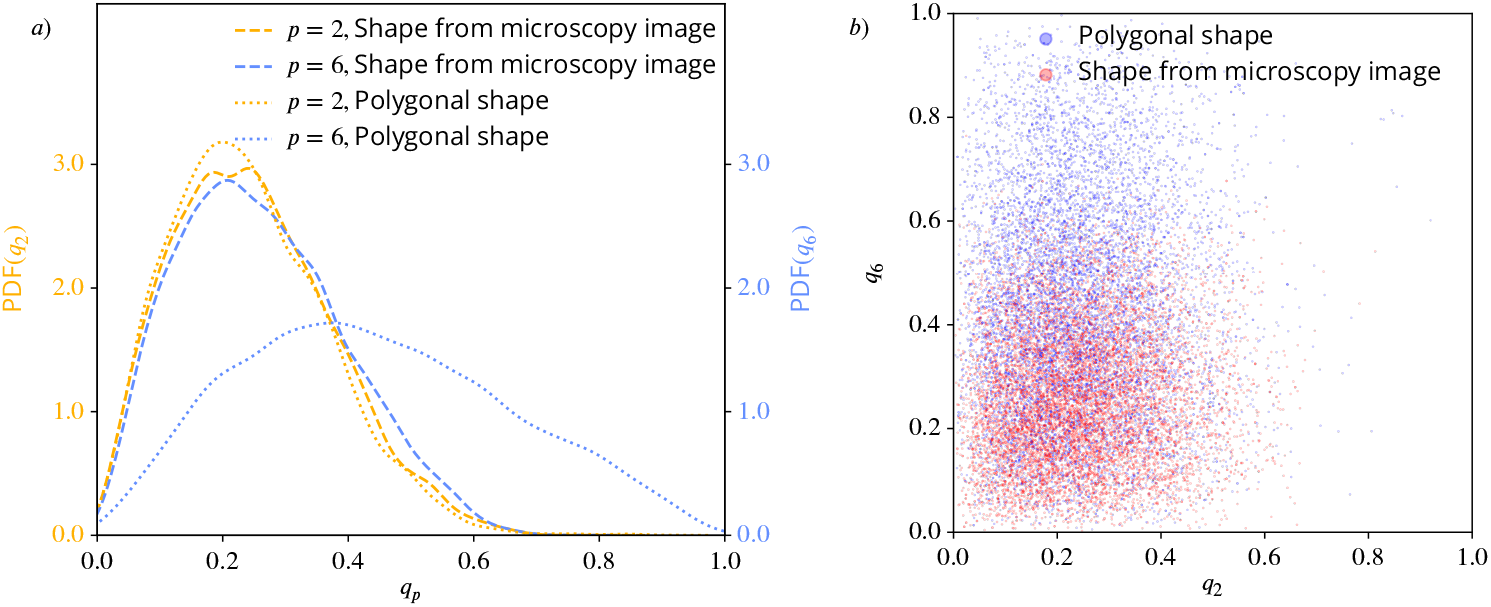
Nematic (*p* = 2) and hexatic (*p* = 6) order for the cells in the experiments from ***Armengol-Collado et al. (2023***).*a*): Probability distribution functions (PDFs) using kde-plots, for *q*_2_ (yellow) and *q*_6_ (blue), once using the polygonal approximation of the cell shape and once using the detailed cell outline obtained from the microscopy pictures. *b*): *q*_6_ (y-axis) versus *q*_2_ (x-axis) for all cells from the experimental data in ***Armengol-Collado et al. (2023***), once using the polygonal approximation of the cell shape (blue) and once using the detailed cell outline obtained from the microscopy pictures (red). For each cell and each timestep we plot one point (*q*_2_, *q*_6_). **Figure 10—figure supplement 1**. Experimental image, segmented cell outline and polygonal shape **Figure 10—figure supplement 2**. Distance correlation *dCor*(*q*_2_, *q*_6_) between *q*_2_ and *q*_6_ **Figure 10—figure supplement 3**. P-values of the distance correlation 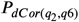

We also test the values for *q*_2_ and *q*_6_ for independence, see ***Figure 10*** b) and in the corresponding statistical measures ***Figure 10—figure Supplement 2*** and ***Figure 10—figure Supplement 3***. The indicated independence in the scatter plots ***Figure 10*** b) is confirmed by the distance correlation and the P-values, as in ***Figure 7, Figure 7—figure Supplement 1*** and ***Figure 7—figure Supplement 2***. This holds for the polygonal shapes as well as for the more detailed shapes from the microscopy images.

As a consequence, the same arguments as discussed above also hold for the experimental data and thus caution against interpretation of *q*_2_ and *q*_6_ or their coarse-grained quantities *Q*_2_ and *Q*_6_ as interdependent order parameters. However, in order to compare with ***Armengol-Collado et al. (2023***) we next compute the averaged coarse-grained quantities 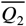 and 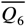 for various coarse-graining radii *R*. In ***Figure 11*** these curves are shown for the polygonal shapes (a)) and the microscopy images (b)). As in ***Armengol-Collado et al. (2023***) we carried out the coarse-graining until the coarse-graining radius corresponds to half of the domain width. In both plots the curves for 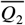 are almost identical, reflecting the similar PDFs in ***Figure 10*** a). For 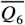 the slope is similar but the curves are shifted. A potential crossover therefore also depends on the approximation of the cells. In any case, for the considered data no consistent hexatic-nematic crossover can be observed.

**Figure 11.**
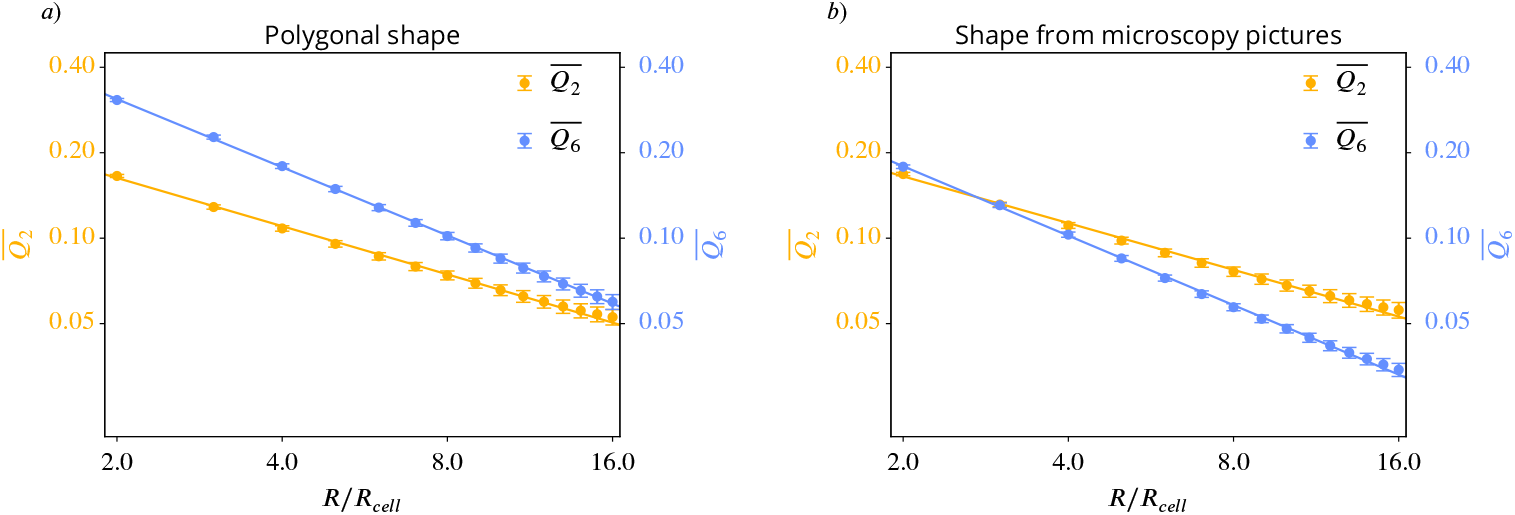
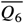 versus 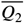 for different coarse graining radii for the experimental data from ***Armengol-Collado et al. (2023***). On the left side *a*) we use the polygonal approximation of the cell shape, on the right side *b*) we use the detailed cell outline obtained from the microscopy pictures. *Q*_*p*_ was calculated according to ***Equation 11***, the averaging of this and the choice of *R*_𝒞_ follow the description in Coarse-grained quantities. The maximal coarse-graining radius corresponds to half the domain width. A logarithmic scaling was used for both axis. Error bars are obtained as s.e.m..

In order to resolve the discrepancy of these results with ***Armengol-Collado et al. (2023***) we next examine the analysis using the alternative shape measures *γ*_*p*_ in ***Equation 8***, which have been considered in ***Armengol-Collado et al. (2023***) but are shown to be not stable.

### Sensitivity of the results on the considered shape descriptor

We now demonstrate that the alternative shape descriptors *γ*_*p*_ in ***Equation 8***, which have been used in (***Armengol-Collado et al., 2023***), can lead to qualitatively different and thus misleading results. The corresponding figures to ***Figure 1, Figure 2*** and ***Figure 4*** are shown in ***Appendix 3 Figure 22, Appendix 3 Figure 23*** and ***Appendix 3 Figure 21***, respectively. For the experimental data in ***Figure 1*** and ***Figure 2*** we use the Voronoi interface method (***Saye and Sethian, 2011, 2012***) to calculate the vertices of a polygon approximating the cell shape. The comparison of these figures already indicates differences between the two methods. Such differences can be seen in the approximated cell shapes, the probability distribution functions (PDFs) and more quantitatively also by comparing 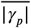 in ***Appendix 3 Figure 23*** *a*) with 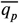 in ***Figure 2*** *a*), which, e.g., for *p* = 6 almost double.

For a more detailed comparison of |*γ*_*p*_| and *q*_*p*_ we investigate the data from the active vertex model and the polygonal approximation of the cells in ***Armengol-Collado et al. (2023***). We restrict ourselves to this data as for multiphase field data the usage of *γ*_*p*_ first requires the approximation of a cell by a polygon and we have already seen that this approximation strongly influences the results.

We focus on ***Figure 8*** and the corresponding results in ***Figure 12***. Instead of the monotonic trend for activity and deformability, which was found for 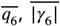 exhibits non-monotonic trends, peaking at intermediate values of activity or deformability. For completeness we also provide the corresponding figures to ***Figure 7*** and ***Figure 9*** in ***Appendix 3 Figure 24*** and ***Appendix 3 Figure 25***, respectively.

**Figure 12.**
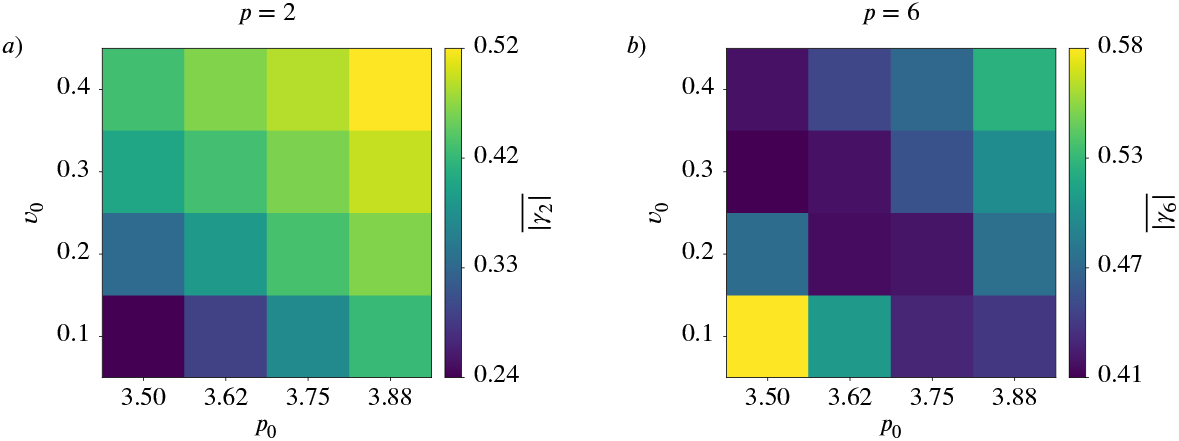
Mean value 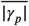 as function of deformability *p*_0_ and activity *v*_0_ for active vertex model. a) nematic order (*p* = 2), b) hexatic order (*p* = 6).

For the polygonal data of the MDCK cells considered in (***Armengol-Collado et al., 2023***), we compare the coarse-grained quantities 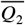 and 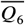, already considered in ***Figure 11*** a), with 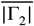 and 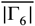 computed from *γ*_2_ and *γ*_6_ using ***Equation 12***, see ***Figure 13***. For completeness we also provide the corresponding figure to ***Figure 10*** in ***Appendix 3 Figure 26***. Besides minor differences regarding the calculation of *R*_*cell*_ ***Figure 13*** corresponds to (***Armengol-Collado et al., 2023***, Fig.3e). Comparing ***Figure 11*** a) and ***Figure 13*** one might come to the conclusion that there is no hexatic-nematic crossover using 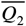 and 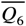, but there is a hexatic-nematic crossover using 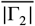 and 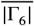. As the only difference between these two evaluations is the considered shape characterization this analysis adds another argument that the proposed crossover is not a robust physical feature of the system.

**Figure 13.**
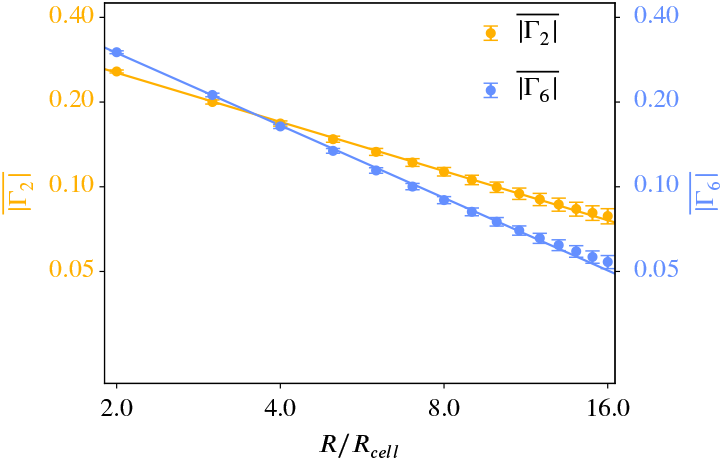
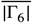 versus 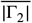 for different coarse graining radii for the experimental data from ***Armengol-Collado et al. (2023***). We use only the polygonal approximation of the cell shape as *γ*_*p*_ can only work with polygons. *Q*_*p*_ was calculated according to ***Equation 12***, the averaging of this and the choice of *R*_𝒞_ follow the description in Coarse-grained quantities. The maximal coarse-graining radius corresponds to half the domain width. A logarithmic scaling was used for both axis. Error bars are obtained as s.e.m..

These results also demonstrate that not only the approximation of the cell boundaries by polygonal shapes heavily influences the characterization of *p*-atic order but also the considered method to classify the shape might lead to qualitative different results. This confirms the argumentation in Methods and materials that Minkowski tensors should be preferred because of their stability properties.

## Discussion

In this study, we introduced Minkowski tensors as a robust and versatile tool for quantifying *p*-atic order in multicellular systems, particularly in scenarios involving rounded or irregular cell shapes. By applying this framework to extensive datasets from two distinct computational models—the active vertex model and the multiphase field model—we identified universal trends: increasing activity and deformability of the cells enhance nematic order (*p* = 2) while diminishing hexatic order (*p* = 6). The consistency of these findings across two models, despite their inherent differences, underscores the generality of our results.

While various shape characterization methods, such as the bond order parameter (***Loewe et al., 2020***; ***Monfared et al., 2023***) and the shape function *γ*_*p*_ (***Armengol-Collado et al., 2023***), have been explored in the literature, we demonstrated that the choice of shape descriptor significantly impacts the conclusions drawn. Such divergences, together with limited mathematical foundations, highlight the limitations of these alternative shape measures in capturing consistent patterns and emphasize the need for stable, reliable shape measures like the Minkowski tensors. As the stability of Minkowski tensors - in contrast to the bond order parameter or the shape function *γ*_*p*_ - can be mathematically justified Minkowski tensors should be the preferred shape descriptor. This finding is not merely a technical nuance, it leads to qualitative differences. Analyzing experimental data for MDCK cells, e.g., has demonstrated that a strong hexatic order on the cellular scale has no physical origin but is a consequence of the approximation of the cell boundaries by polygonal shapes, which is a requirement to use the shape function *γ*_6_. Considering the full cellular boundaries and *q*_6_, which is derived from the Minkowski tensors, leads to a different picture, with weaker hexatic order. A critical question in the literature has been whether shape measures for different *p*-atic orders can be directly compared. While some studies have suggested relationships between *γ*_2_ and *γ*_6_ (***Armengol-Collado et al., 2023***), our results refute this notion. We demonstrated that measures like *q*_2_ and *q*_6_ are independent and capture fundamentally distinct aspects of cell shape and alignment. Comparing them directly is mathematically but also physically and biologically misleading. We further tested the hypothesis of a hexatic-nematic crossover at larger length scales by coarse-graining *q*_2_ and *q*_6_. To discuss such a crossover requires direct comparison of *q*_2_ and *q*_6_ or their coarse-grain quantities *Q*_2_ and *Q*_6_, which is conceptually questionable. However, the results showed no consistent trends indicative of a crossover, regardless of the considered model or the experimental data. This leads to the conclusion that the proposed hexatic-nematic crossover in (***Armengol-Collado et al., 2023***) is not a physical phenomena but specific to the considered method.

Our findings suggest that *p*-atic orders should be studied independently, also across length scales, as they describe complementary aspects of cellular organization. The coexistence of distinct orientational orders emphasized in different studies—such as nematic (*p* = 2) (***Duclos et al., 2017***; ***Saw et al., 2017***; ***Kawaguchi et al., 2017***), tetratic (*p* = 4) (***Cislo et al., 2023***), and hexatic (*p* = 6) (***Li and Ciamarra, 2018***)—is not contradictory but highlights the rich, multifaceted nature of cellular organization. Rather than searching for a single dominant order, future research should focus on the interplay of different *p*-atic orders and their associated defects. This suggests to not only consider *p*-atic liquid crystal theories (***Giomi et al., 2022a***) for one specific *p*, but combinations of these models for various *p*’s. Understanding how these orders interact may reveal how they collectively regulate morphogenetic processes.

Connecting *p*-atic orders to biological function remains a critical avenue for exploration. While the mathematical independence of *q*_2_, *q*_6_, and other shape measures precludes the identification of a universal dominant order, biological systems may exhibit context-dependent preferences. For example, a specific *p*-atic order might correlate with or drive a key morphogenetic event. Investigating these connections could yield insights into how tissues achieve functional organization and adapt to environmental cues. As such, while it is difficult to speak of dominating orders from a mathematical point of view, there could be a dominating order from a biological point of view, meaning the *p*-atic order connected to the governing biological process.

## Data availability

A code illustrating the extraction of the contour and the calculation from *q*_*p*_ and *ϑ*_*p*_ for grayscale images can be found on Zenodo at https://doi.org/10.5281/zenodo.15430268.

## Acknowledgments

We acknowledge fruitful discussions with Björn Böttcher, Brendan Tobin and Emma Happel. HJ acknowledges funding by the European Union’s Horizon 2020 research and innovation programme under the Marie Skłodowska-Curie grant agreement No. 945371. RS acknowledges support from the UK Engineering and Physical Sciences Research Council (Award EP/W023946/1). AD acknowledges funding from the Novo Nordisk Foundation (grant No. NNF18SA0035142 and NERD grant No. NNF21OC0068687), Villum Fonden (grant No. 29476), and the European Union (ERC, PhysCoMeT, 101041418). AV acknowledges funding from the German Research Foundation (Award FOR3013 “Vector- and tensor-valued surface PDEs”) and computing resources provided by JSC through MORPH and by ZIH through WIR. Views and opinions expressed are however those of the authors only and do not necessarily reflect those of the European Union or the European Research Council. Neither the European Union nor the granting authority can be held responsible for them.

## Appendix 1

### Parameters for the computational models

**Appendix 1—table 1.** Values of the dimensionless parameters used in the active vertex model.

| Parameter | Description | Numerical Value |
| --- | --- | --- |
| $N$ | Number of cells | 100 |
| $L$ | Simulation box size | 10 |
| $T$ | Total simulation time | 300 |
| $k_p$ | Perimeter elastic modulus | 1.0 |
| $\tau$ | Simulation time step | 0.01 |
| $v_0$ | Self-propulsion strength (i.e. activity) | 0.1 - 0.4 |
| $D_r$ | Rotation diffusion coefficient | 0.05 |
| $p_0$ | Shape index of active cells | 3.5 - 3.875 |

**Appendix 1—table 2.** Values of the dimensionless parameters used in the multiphase-field model simulations.

| Parameter | Description | Numerical Value |
| --- | --- | --- |
| $N$ | Number of cells | 100 |
| $L$ | Simulation box size | 100 |
| $T$ | Total simulation time | 150 |
| $\epsilon$ | Interface width | 0.15 |
| $\tau$ | Simulation time step | 0.005 |
| $v_0$ | Self-propulsion strength (i.e. activity) | 0.4 - 1.0 |
| $D_r$ | Rotation diffusion coefficient | 0.01 |
| $\alpha$ | Alignment parameter | 0.1 |
| $Ca$ | Capillary number | 0.05 - 0.2 |
| $In$ | Interaction number | 0.1 |
| $a_a$ | Cell-cell attraction strength | 1.0 |
| $a_r$ | Cell-cell repulsion strength | 1.0 |
| $h$ | Mesh size | $5h \approx \epsilon$ |

## Appendix 2

### Results for *q*_3_,*q*_4_ and *q*_5_

#### Distribution functions in dependence of deformability

**Appendix 2—figure 14.**
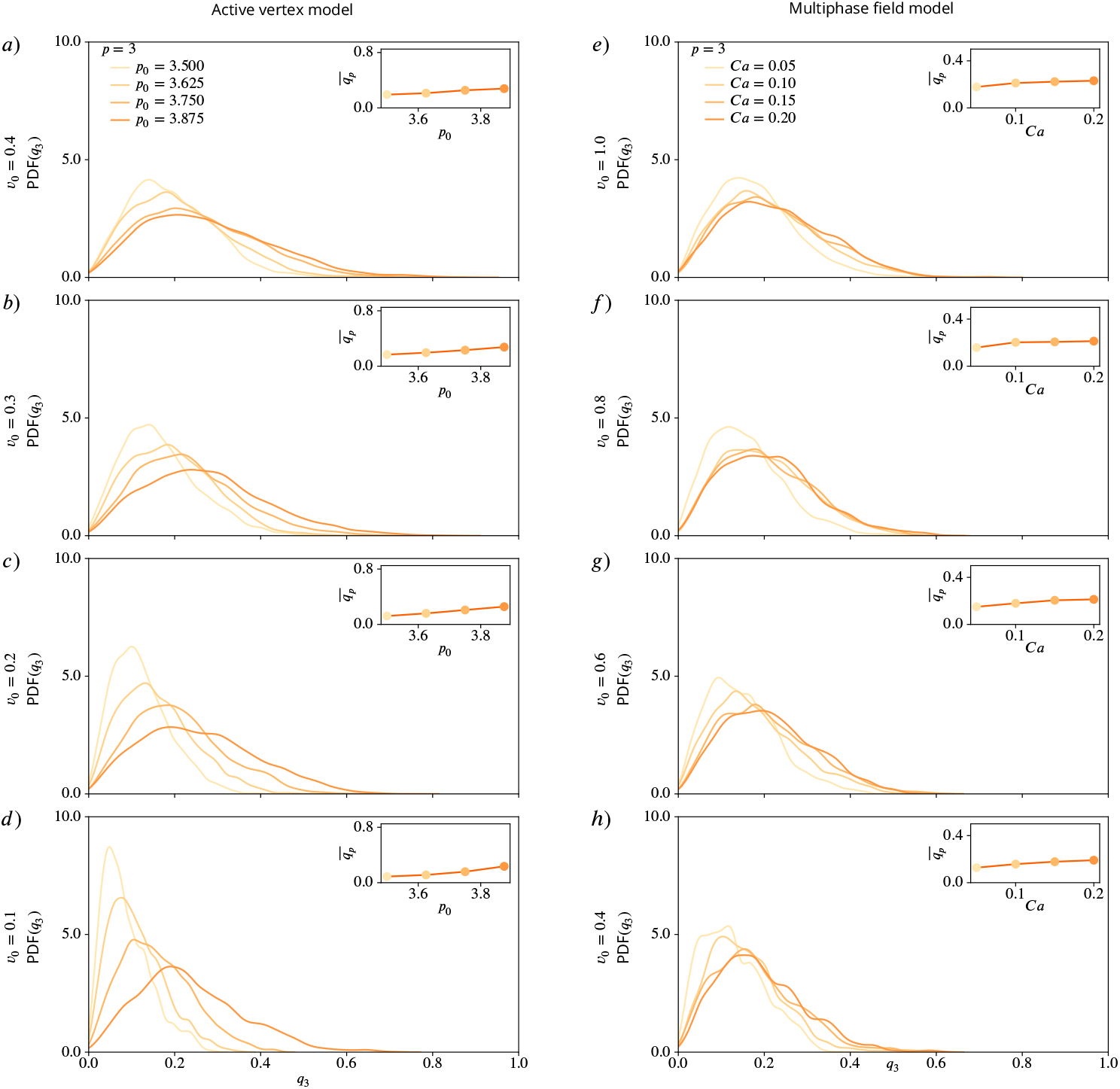
PDFs for *q*_3_ using kde-plots, for varying deformability *p*_0_ or *Ca* and fixed activity *v*_0_. Inlets show mean values of *q*_3_ as function of deformability. a) - d) active vertex model, e) - h) multiphase field model for decreasing activity.

**Appendix 2—figure 15.**
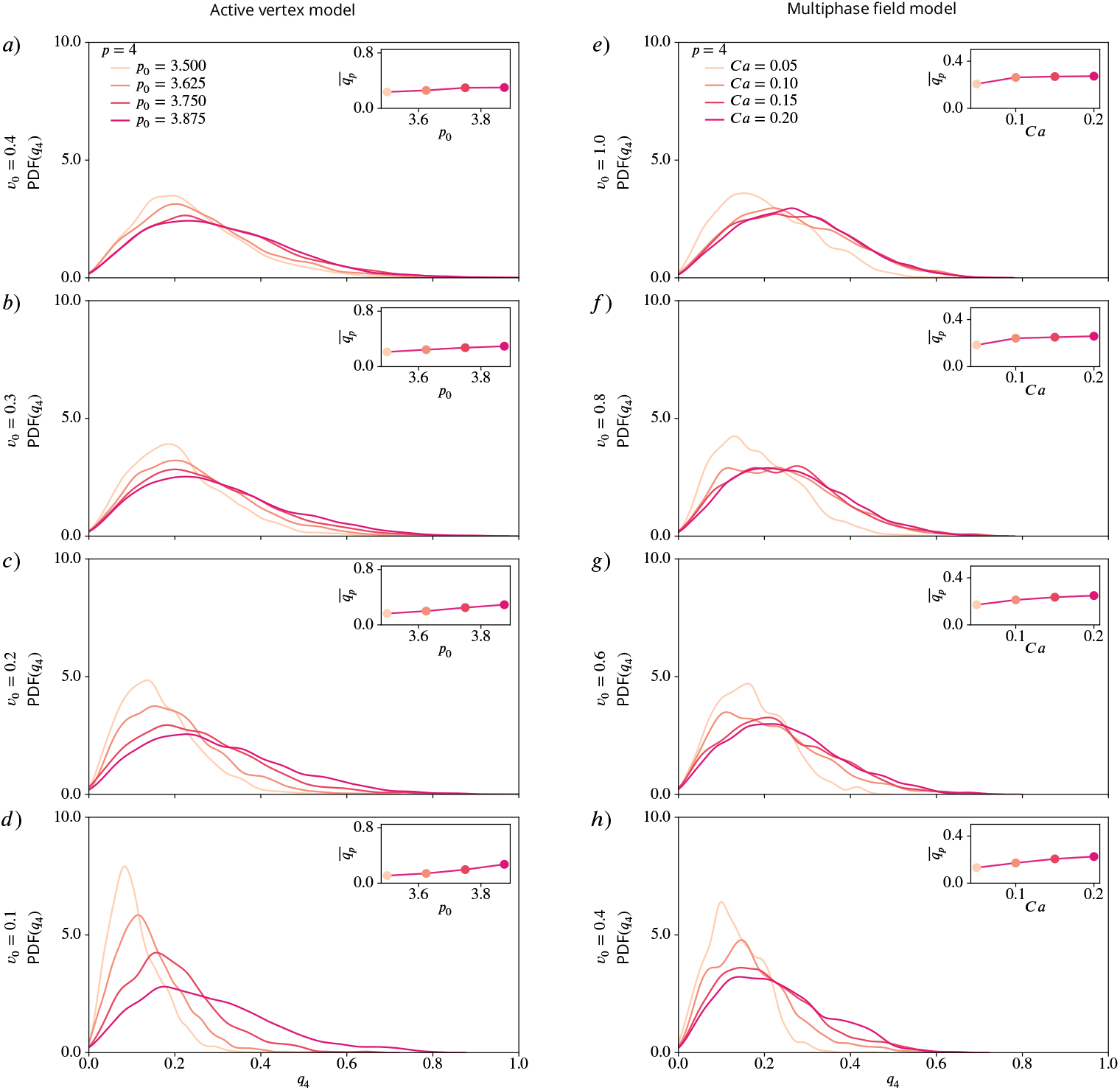
PDFs for *q*_4_ using kde-plots, for varying deformability *p*_0_ or *Ca* and fixed activity *v*_0_. Inlets show mean values of *q*_4_ as function of deformability. a) - d) active vertex model, e) - h) multiphase field model for decreasing activity.

**Appendix 2—figure 16.**
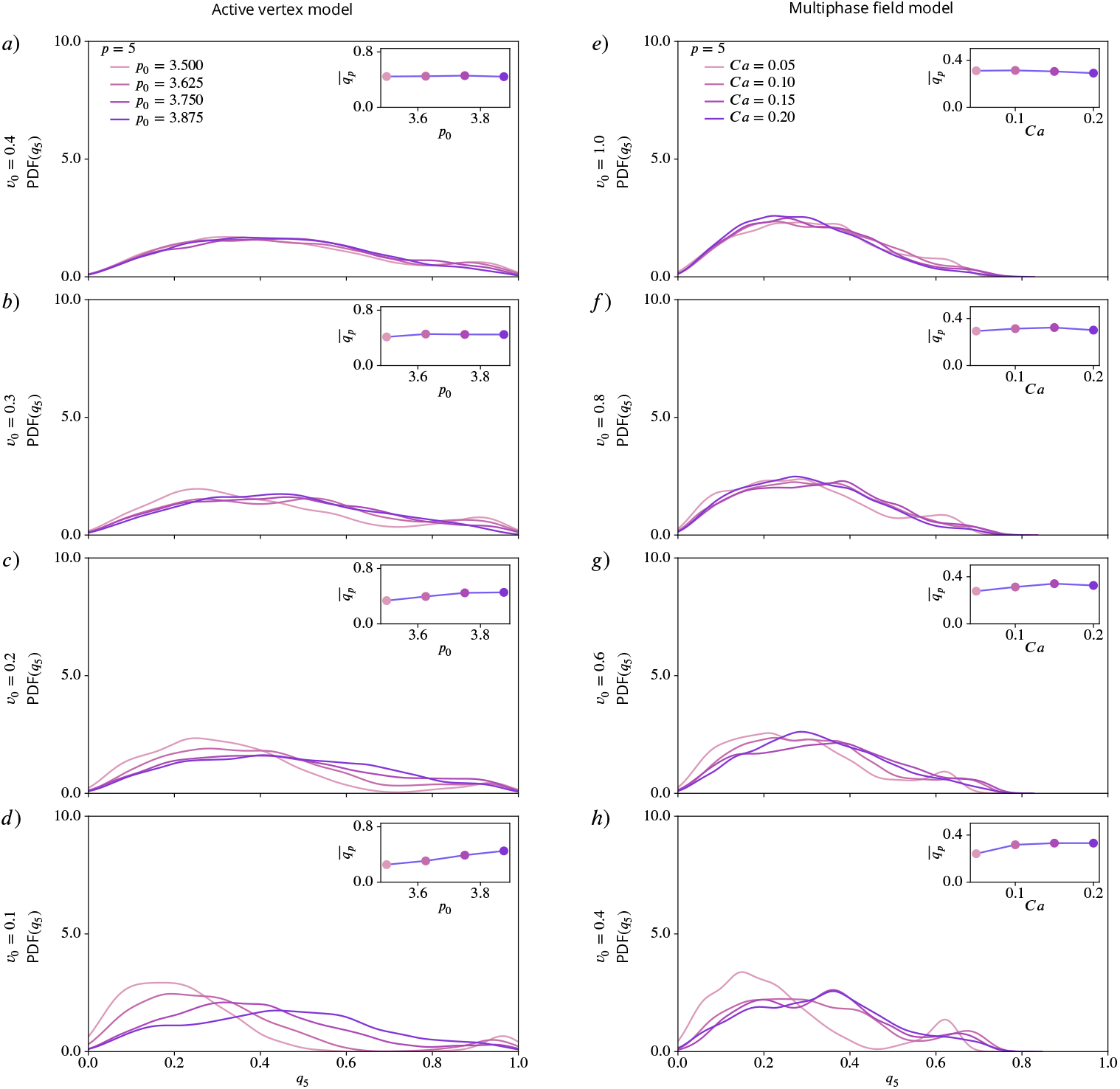
PDFs for *q*_5_ using kde-plots, for varying deformability *p*_0_ or *Ca* and fixed activity *v*_0_. Inlets show mean values of *q*_5_ as function of deformability. a) - d) active vertex model, e) - h) multiphase field model for decreasing activity.

#### Distribution functions in dependence of activity

**Appendix 2—figure 17.**
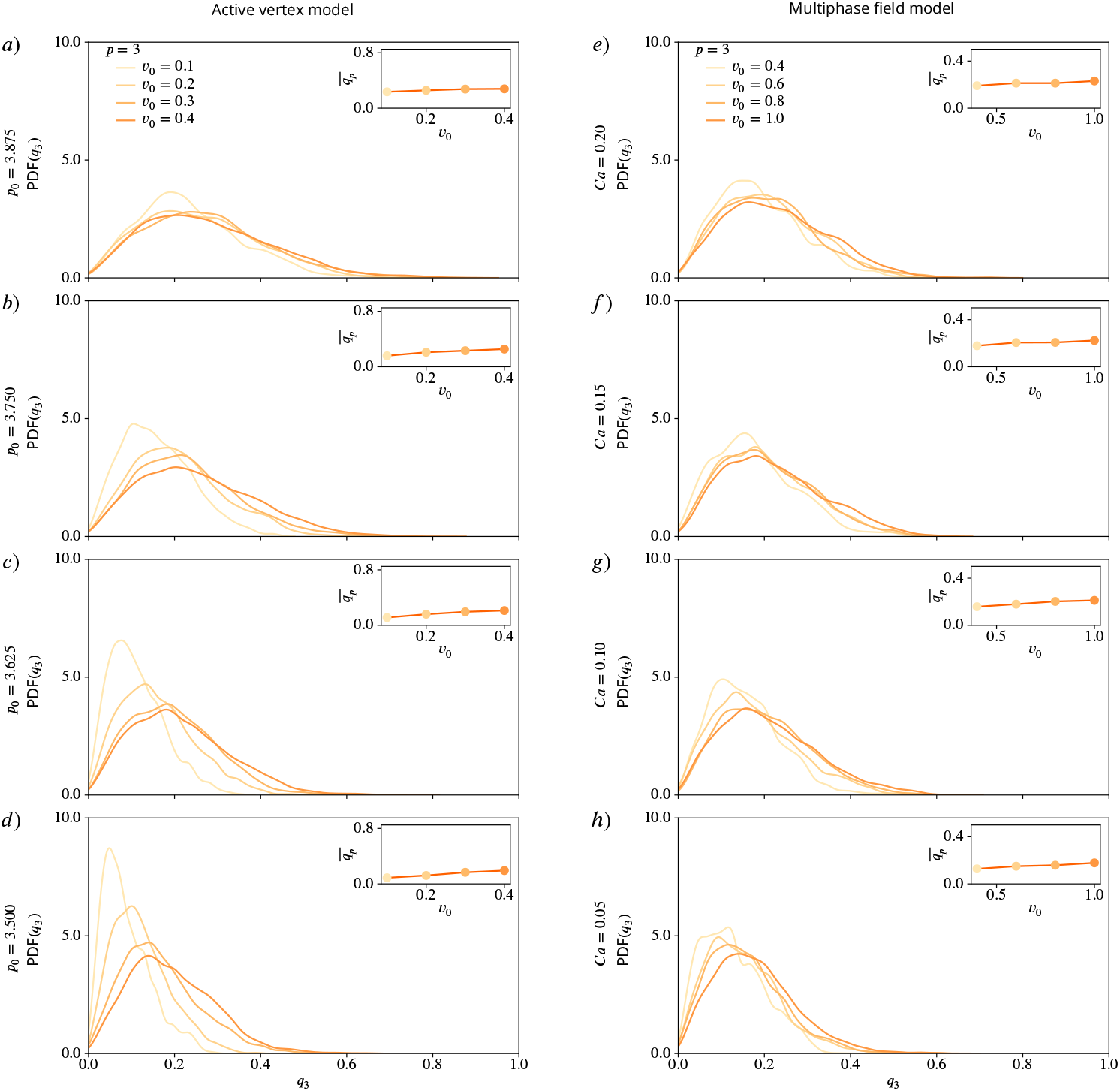
PDFs for *q*_3_ using kde-plots, for varying activity *v*_0_ and fixed deformability *p*_0_ or *Ca*. Inlets show mean values of *q*_3_ as function of activity. a) - d) active vertex model and e) - h) multiphase field model for decreasing deformability.

**Appendix 2—figure 18.**
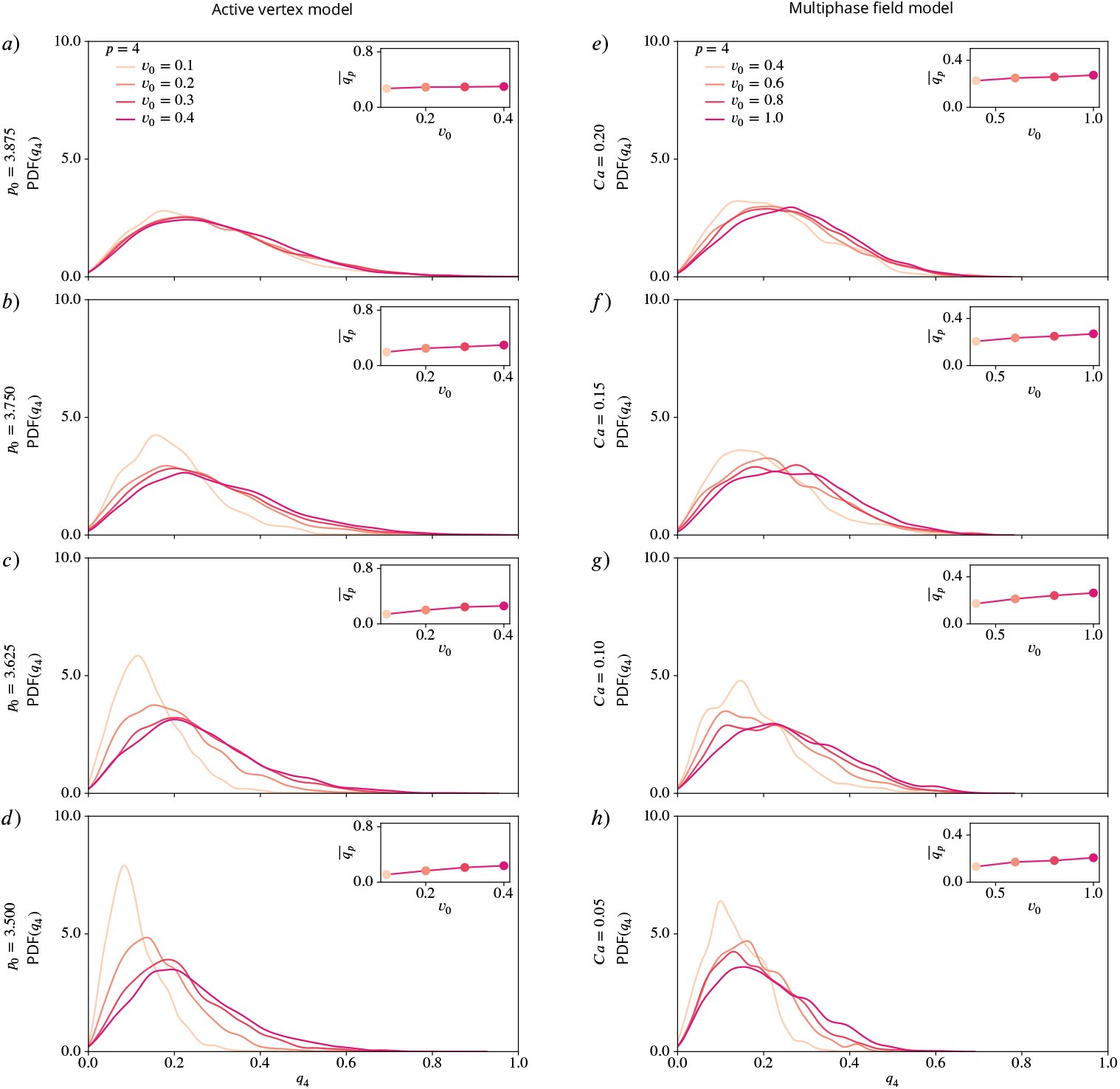
PDFs for *q*_4_ using kde-plots, for varying activity *v*_0_ and fixed deformability *p*_0_ or *Ca*. Inlets show mean values of *q*_4_ as function of activity. a) - d) active vertex model and e) - h) multiphase field model for decreasing deformability.

**Appendix 2—figure 19.**
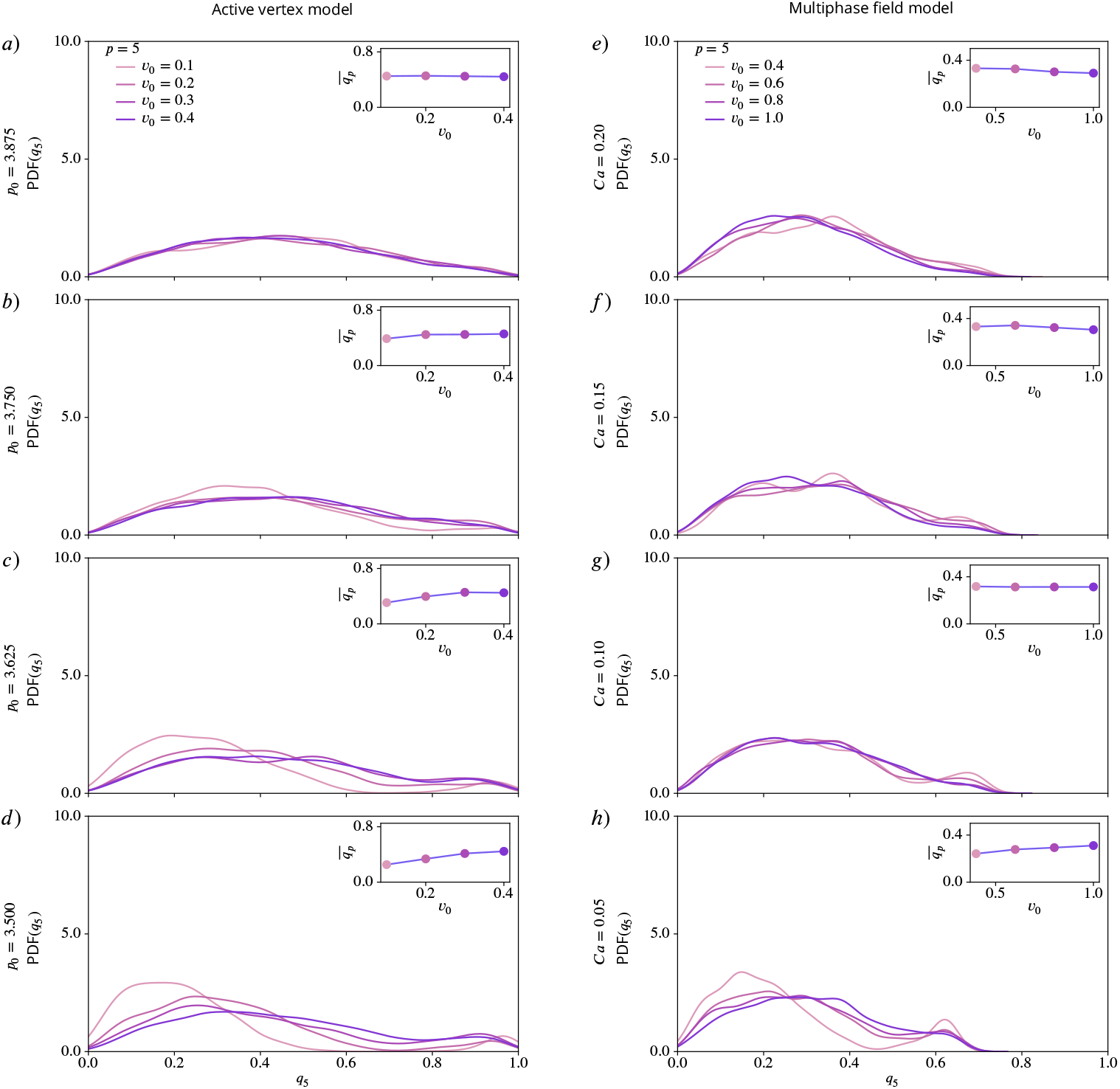
PDFs for *q*_5_ using kde-plots, for varying activity *v*_0_ and fixed deformability *p*_0_ or *Ca*. Inlets show mean values of *q*_5_ as function of activity. a) - d) active vertex model and e) - h) multiphase field model for decreasing deformability.

#### Summarized behavior in dependence of activity and deformability

**Appendix 2—figure 20.**
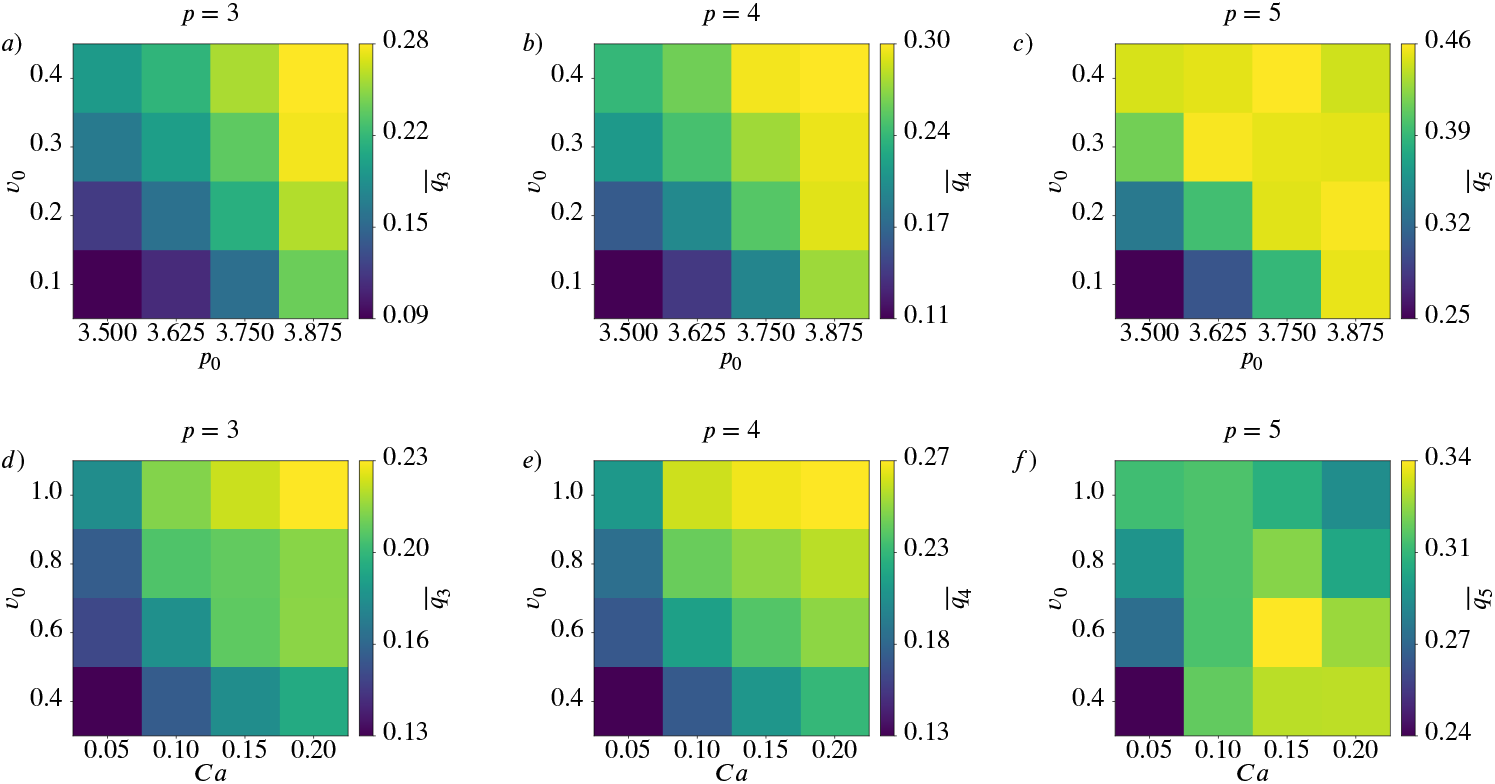
Mean value 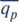 for *p* = 3 (left), *p* = 4 (middle) and *p* = 5 (right) as function of deformability *p*_0_ or *Ca* and activity *v*_0_ for active vertex model (a) - c)) and multiphase field model (d) - f)).

## Appendix 3

### Results using polygonal shape analysis

To visualize the similarities and differences between *q*_*p*_ and *γ*_*p*_ we show the analogue of ***Figure 4*** using the Minkowski tensors in ***Appendix 3 Figure 21*** using polygonal shape analysis. For regular polygons, both shape characterization methods behave in the same way as seen in the first row. For irregular shapes, like in the second row, they already show a different behaviour. Note that the rounded shapes from ***Figure 4*** are missing in ***Appendix 3 Figure 21*** as *γ*_*p*_ requires a polygon.

**Appendix 3—figure 21.**
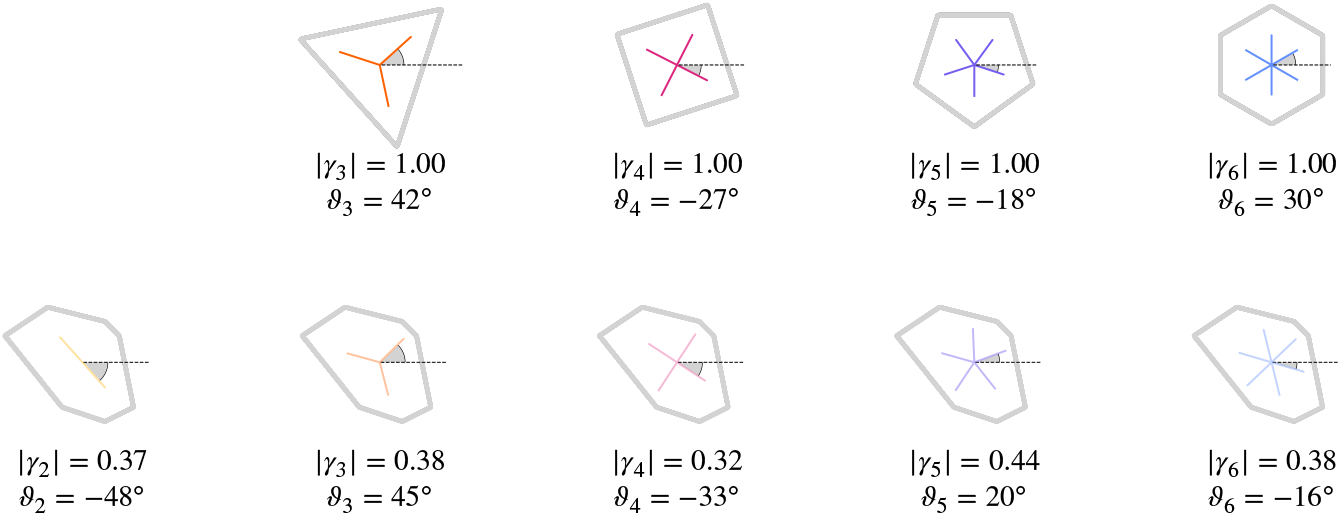
Regular and irregular shapes, adapted from ***Armengol-Collado et al. (2023***), with magnitude and orientation calculated by ***Equation 8*** and ***Equation 9***. The brightness scales with the magnitude |*γ*_*p*_|.

The influence of the choice of the shape characterization method is not only visible in the values for single shapes, it can also be seen in the mean and standard deviation. To illustrate this, we use *γ*_*p*_ to characterize the shapes from the same experimental data as in ***Figure 1*** and ***Figure 2***. The segmented data consists of smooth contours for each cell. As *γ*_*p*_ only works with vertex coordinates the first step is to identify these. Note that this always leads to the approximation of the cell shape with a polygon and therefore to a simplification of the shape. For finding the vertices we consider the Voronoi interface method (***Saye and Sethian, 2011, 2012***). Using the so obtained vertices and ***Equation 8*** and ***Equation 9*** leads to the results shown in ***Appendix 3 Figure 22*** and ***Appendix 3 Figure 23***. Compared to ***Figure 1*** and ***Figure 2***, where the Minkowski tensor was used for the same data, we see that the choice of the shape characterization method highly influences the results.

Qualitative consequences of the approach become apparent by comparing ***Figure 8*** with ***Appendix 3 Figure 12***, which, instead of the monotonic trend for activity and deformability, exhibit nonmonotonic trends, peaking at intermediate values of activity or deformability.

Considering the independence of *γ*_2_ and *γ*_6_ the corresponding results to ***Figure 7*** and the corresponding distance correlation and statistical tests ***Figure 7—figure Supplement 1*** and ***Figure 7— figure Supplement 2***, respectively, are provided in ***Appendix 3 Figure 24*** and ***Appendix 3 Figure 24— figure Supplement 1*** and ***Appendix 3 Figure 24—figure Supplement 24***, respectively, but only for the active vertex model.

**Appendix 3—figure 22.**
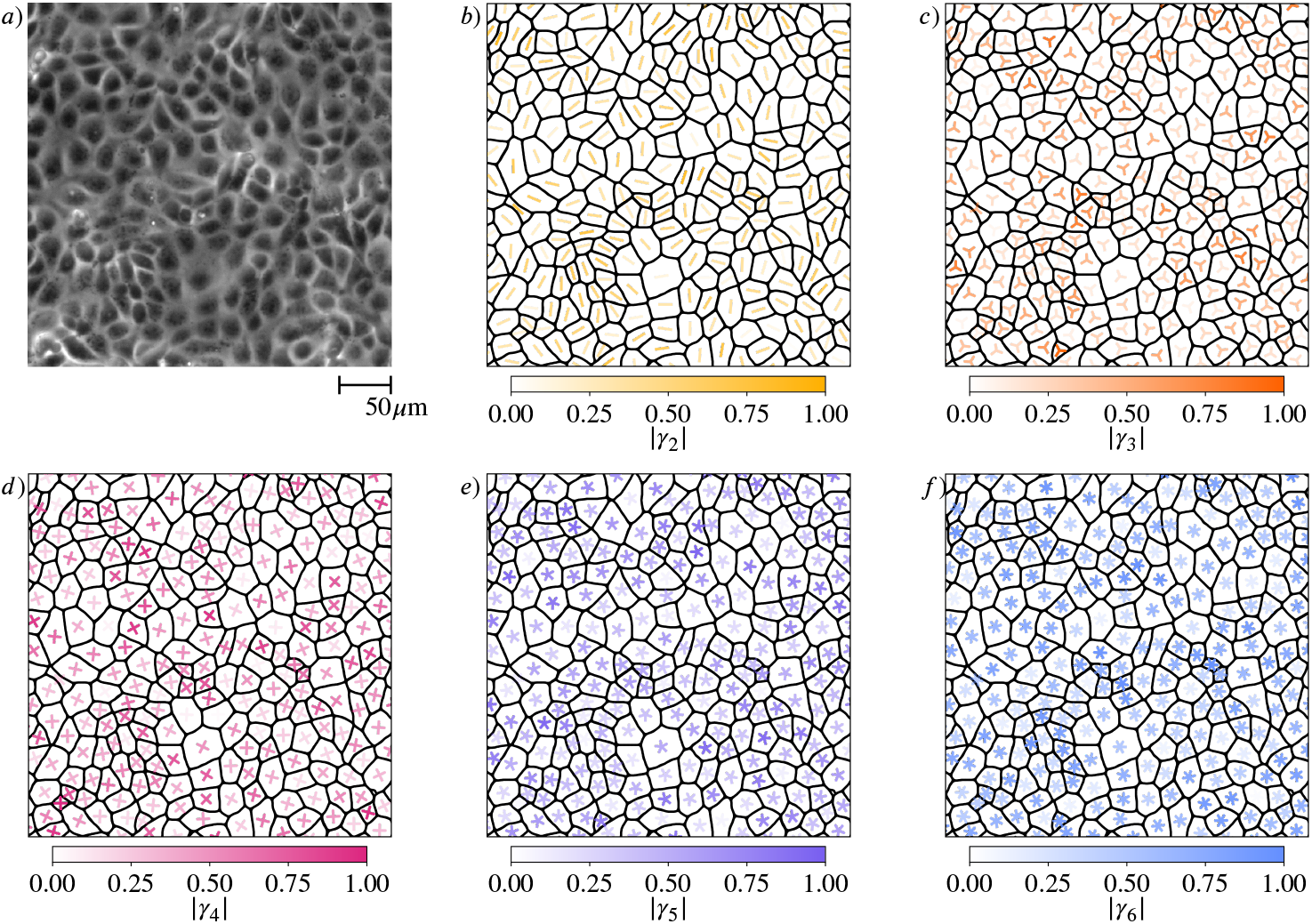
Shape classification of cells in wild-type MDCK cell monolayer. a) Raw experimental data. b) - f) Polygonal shape classification, visualized using *γ*_*p*_ calculated by ***Equation 8*** and ***Equation 9*** for *p* = 2, 3, 4, 5, 6, respectively. The brightness and the rotation of the *p*-atic director indicates the magnitude and the orientation, respectively.

**Appendix 3—figure 23.**
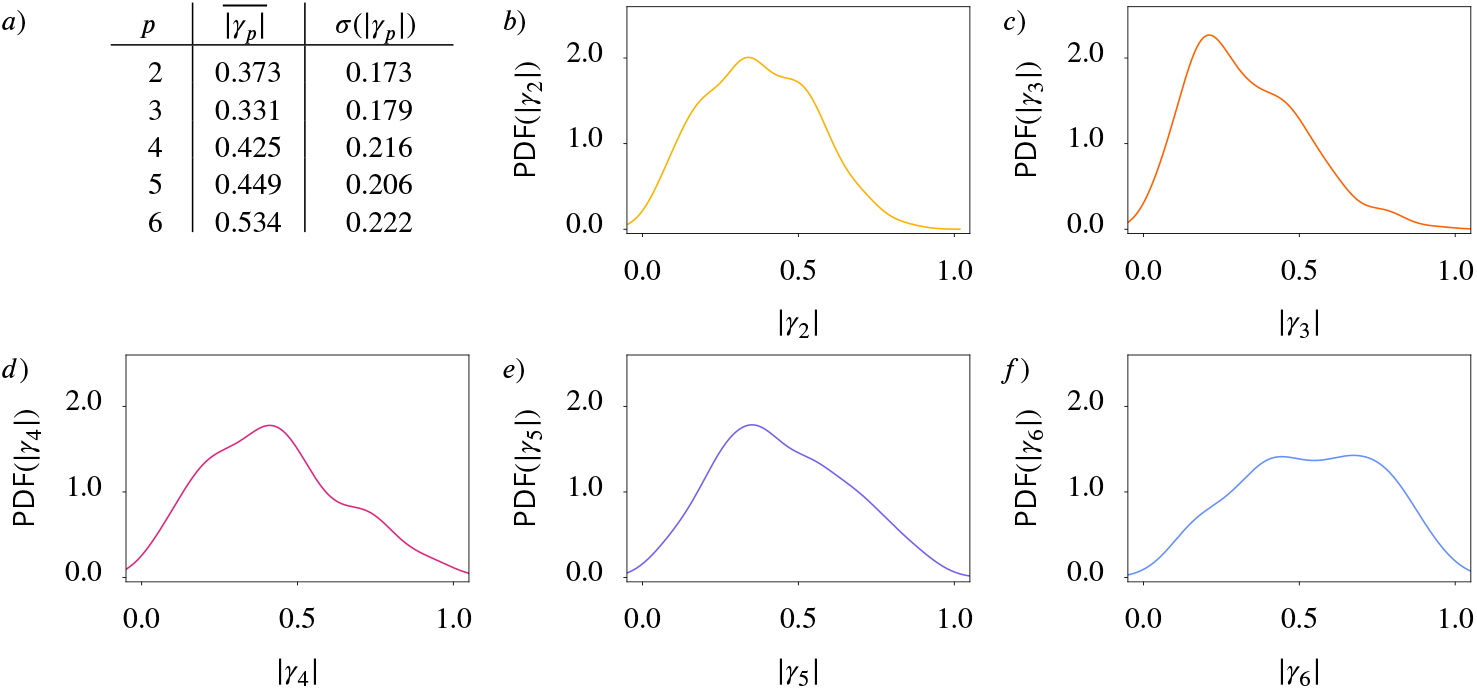
Statistical data for cell shapes identified in ***Appendix 3 Figure 22*** (. a) Mean 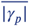 and standard deviation *σ*(| *γ*_*p*_|) of |*γ*_*p*_|. b) - f) Probability distribution function (PDF) of |*γ*_*p*_| for *p* = 2, 3, 4, 5, 6, respectively. Kde-plots are used to show the probability distribution. For this first analysis we regard only one frame with 235 cells.

**Appendix 3—figure 24.**
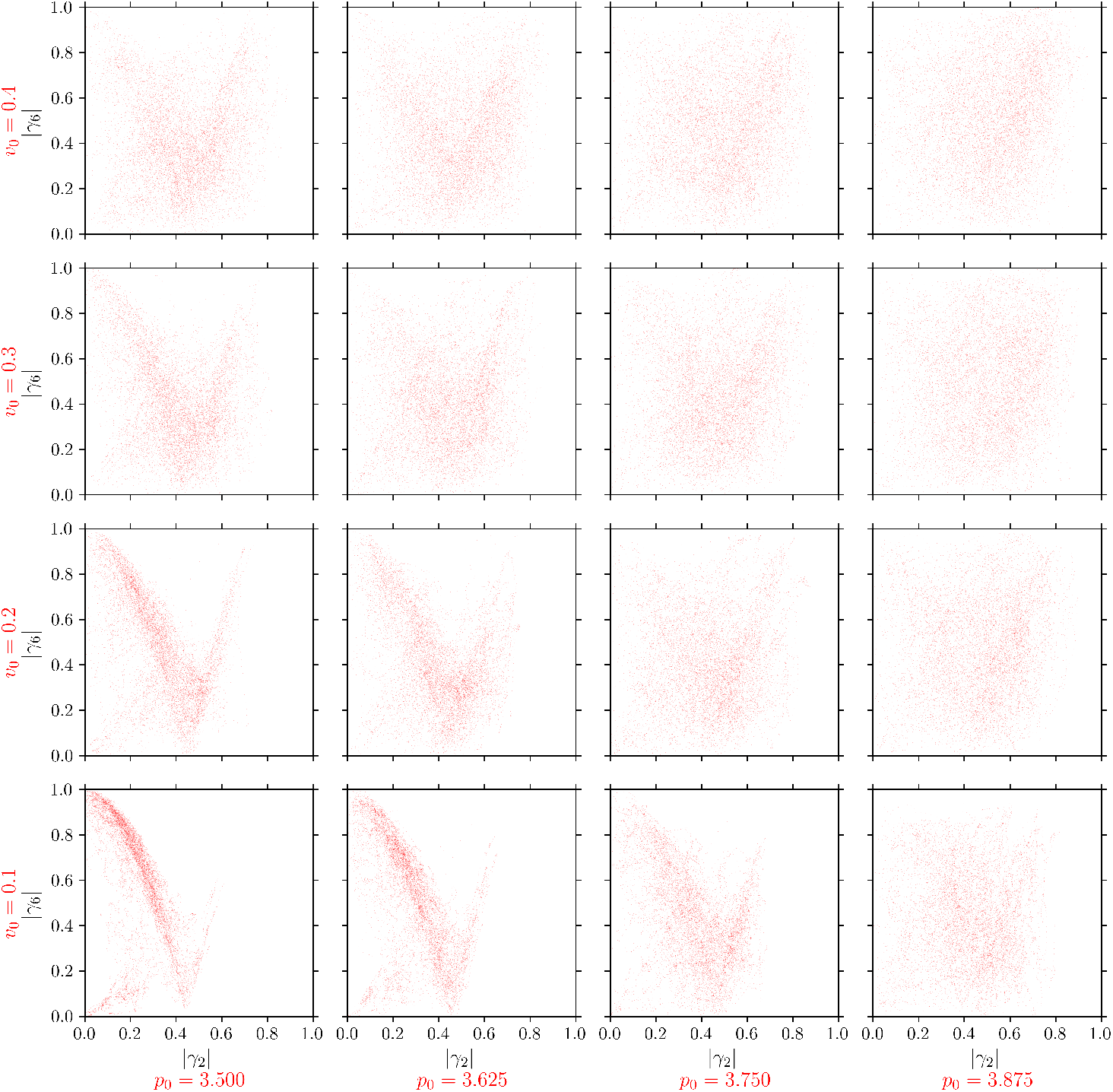
|*γ*_6_| (y-axis) versus |*γ*_2_| (x-axis) for all cells in the active vertex model. For each cell and each timestep we plot one point (|*γ*_2_|, |*γ*_6_|). Each panel corresponds to specific model parameters *p*_0_ and *v*_0_, representing deformability and activity. **Appendix 3—figure 24—figure supplement 1**. Distance correlation *dCor*(| *γ*_2_|, |*γ*_6_|) **Appendix 3—figure 24—figure supplement 2**. P-values of the distance correlation 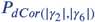

**Appendix 3—figure 25.**
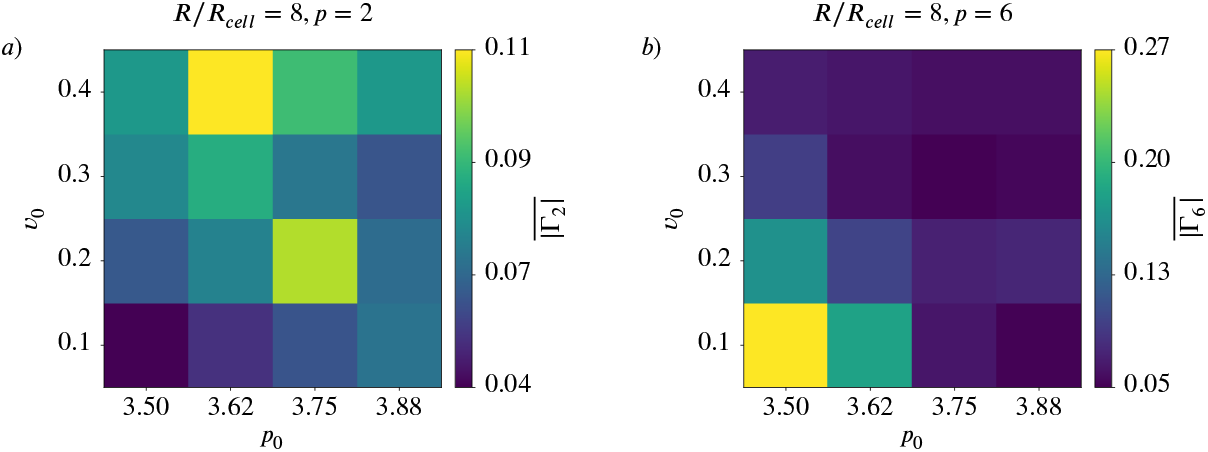
Coarse-Grained nematic (*p* = 2) and hexatic (*p* = 6) order for *R*/*R*_*cell*_ = 8 depending on activity and deformability of the cells. 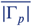 as function of deformability *p*_0_ and activity *v*_0_ for active vertex model. a) nematic order (*p* = 2), b) hexatic order (*p* = 6). **Appendix 3—figure 25—figure supplement 1**. 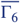 versus 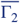 for different coarse graining radii in the active vertex model.

**Appendix 3—figure 26.**
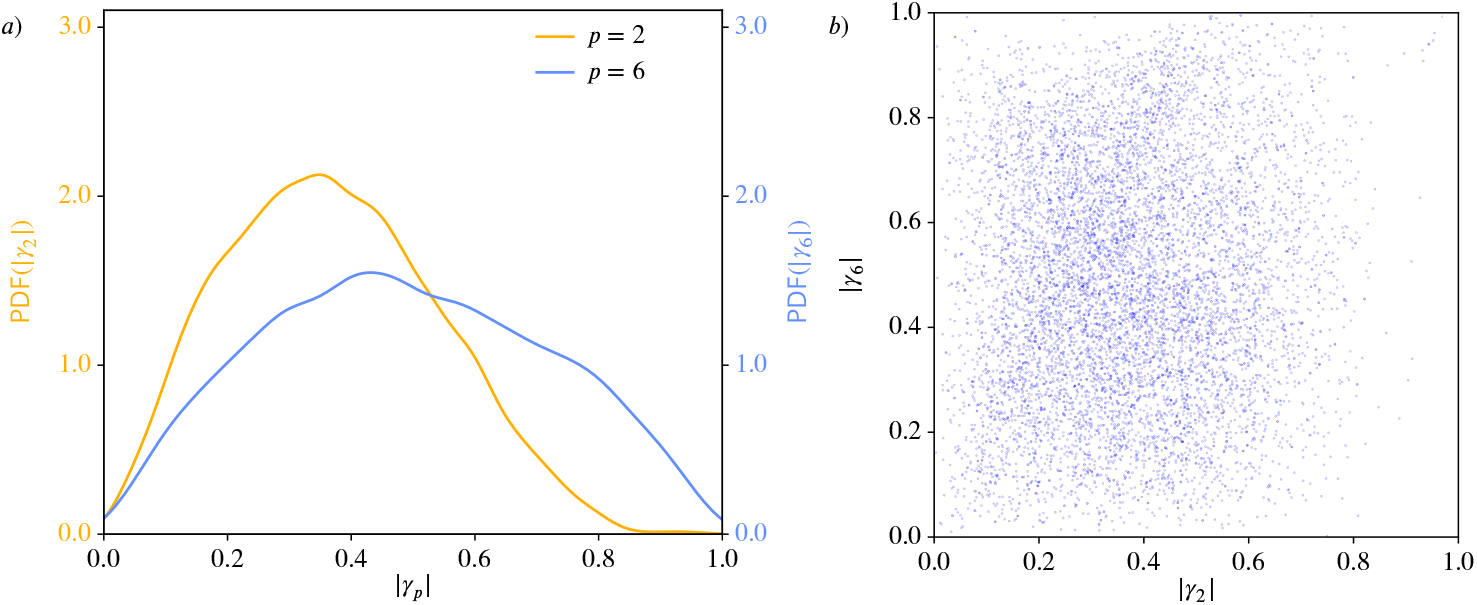
Nematic (*p* = 2) and hexatic (*p* = 6) order for the cells in the experiments from ***Armengol-Collado et al. (2023***). We use the polygonal approximation of the cell shape as *γ*_*p*_ can only work with polygons. *a*): Probability distribution functions (PDFs) using kde-plots, for |*γ*_2_| (orange) and |*γ*_6_| (blue). *b*): Nematic (*p* = 2) and hexatic (*p* = 6) order are independent of eachother. |*γ*_6_| (y-axis) versus |*γ*_2_| (x-axis) for all cells. For each cell and each timestep we plot one point (|*γ*_2_|, |*γ*_6_|). **Appendix 3—figure 26—figure supplement 1**. Distance correlation *dCor*(|*γ*_2_|, |*γ*_6_|) **Appendix 3—figure 26—figure supplement 2**. P-value of the distance correlation 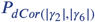

**Figure 7—figure supplement 1.**
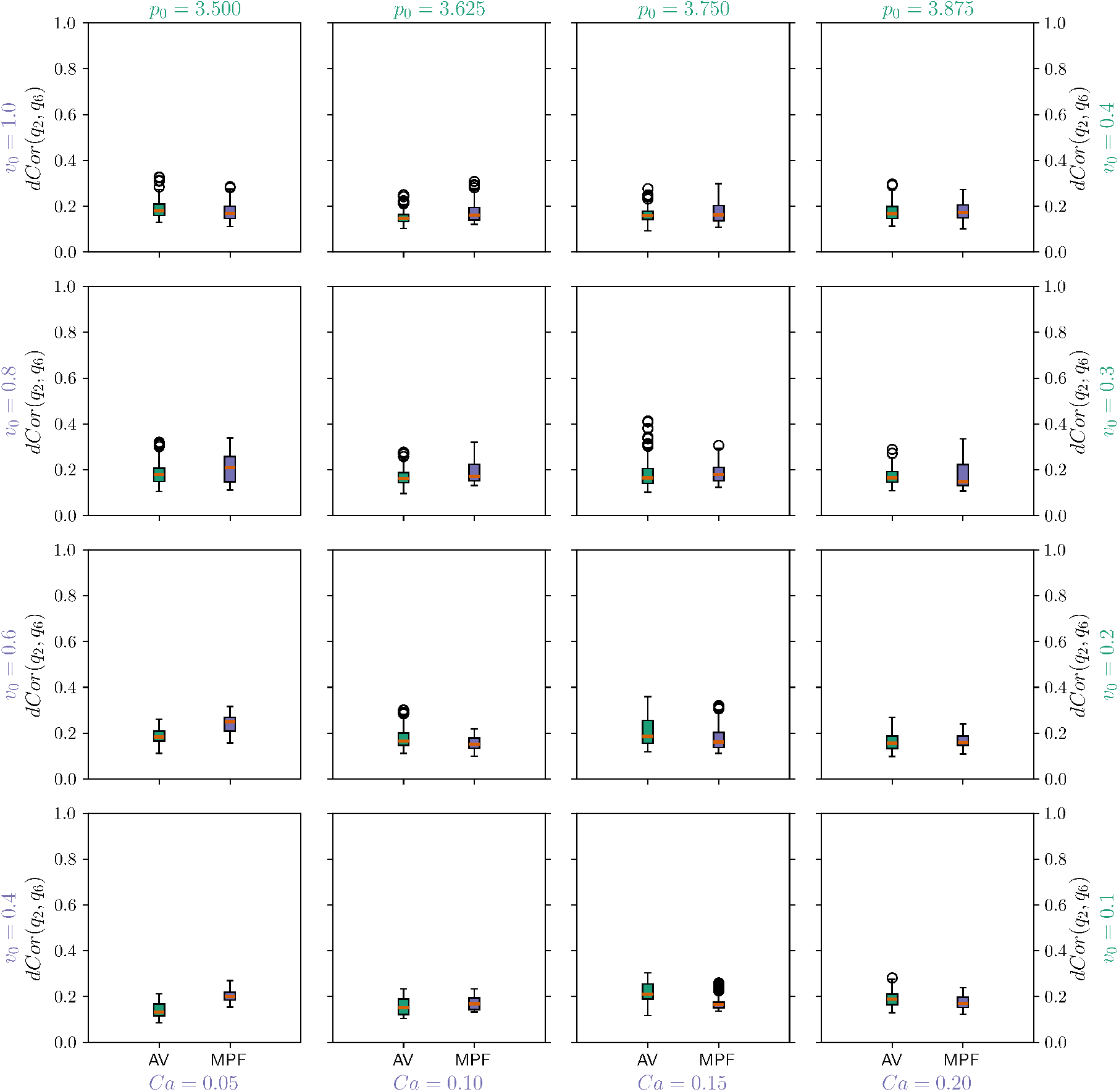
Distance correlation *dCor*(*q*_2_, *q*6) between *q*_2_ and *q*_6_ for all cells in the multiphase field model (MPF - purple box) and active vertex model (AV - green box). Each panel corresponds to specific model parameters; *Ca* and *v*_0_ for multiphase field model, and *p*_0_ and *v*_0_ for the active vertex model, representing deformability and activity, respectively. We compute one value for every timestep and present the resulting distribution with a box plot. In the box plots the orange line corresponds to the median of the data and the box ranges from the first to the third quartile of the data. The whiskers go from the lowest data point greater than (value of the first quartile) − 1.5 × (Interquartile range) to the highest data point below (value of the third quartile) + 1.5 × (Interquartile range). Outliers are shown with circles.

**Figure 7—figure supplement 2.**
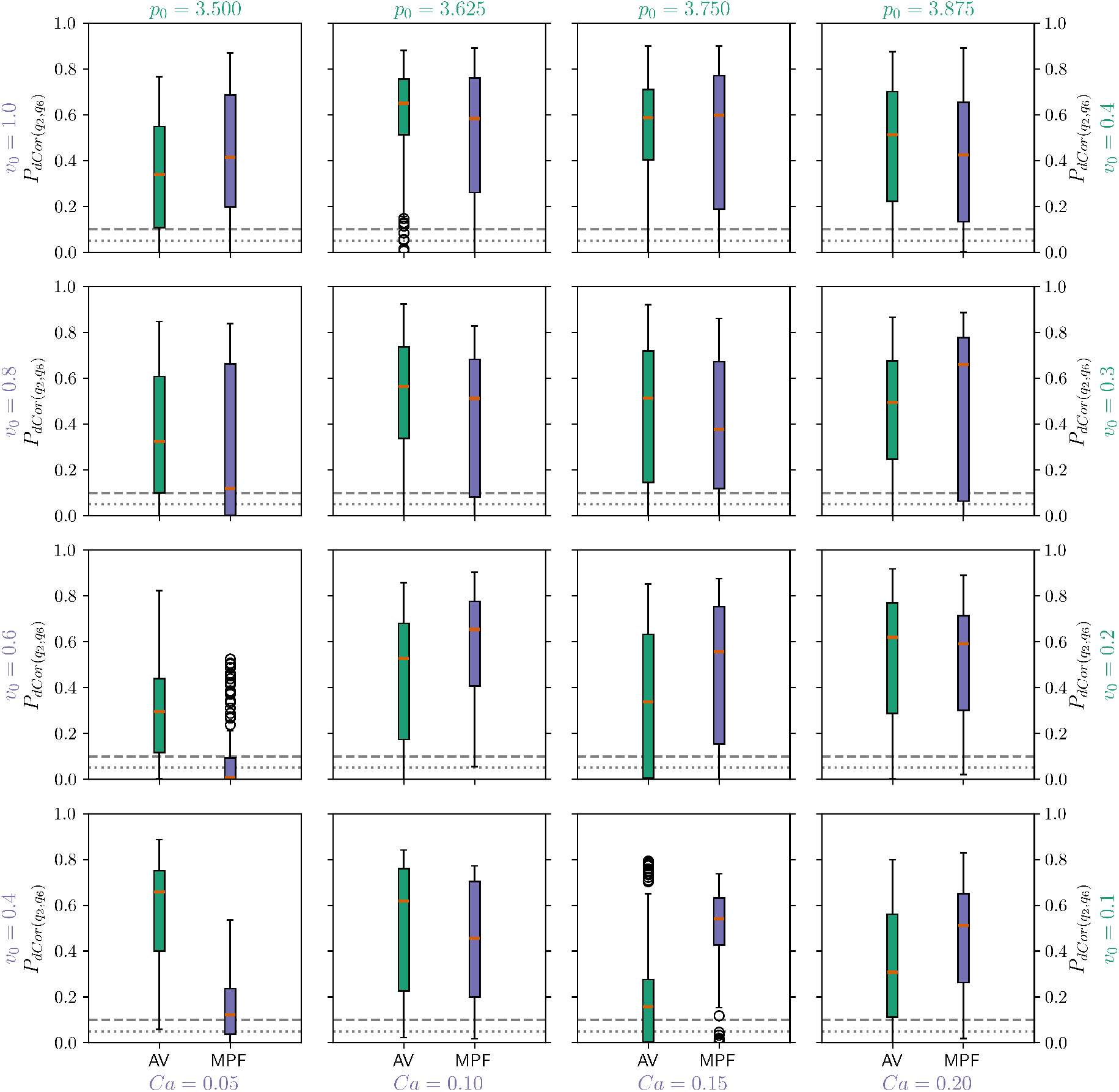
P-values of the distance correlation 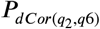 between *q*_2_ and *q*_6_ for all cells in the multiphase field model (MPF - purple box) and active vertex model (AV - green box). Each panel corresponds to specific model parameters; *Ca* and *v*_0_ for multiphase field model, and *p*_0_ and *v*_0_ for the active vertex model, representing deformability and activity, respectively. We compute one value for every timestep and present the resulting distribution with a box plot. In the box plots the orange line corresponds to the median of the data and the box ranges from the first to the third quartile of the data. The whiskers go from the lowest data point greater than (value of the first quartile) − 1.5 ×(Interquartile range) to the highest data point below (value of the third quartile) + 1.5 × (Interquartile range). Outliers are shown with circles. 0.05 and 0.1 are marked with a grey dotted/grey dashed line to guide the eye.

**Figure 8—figure supplement 1.**
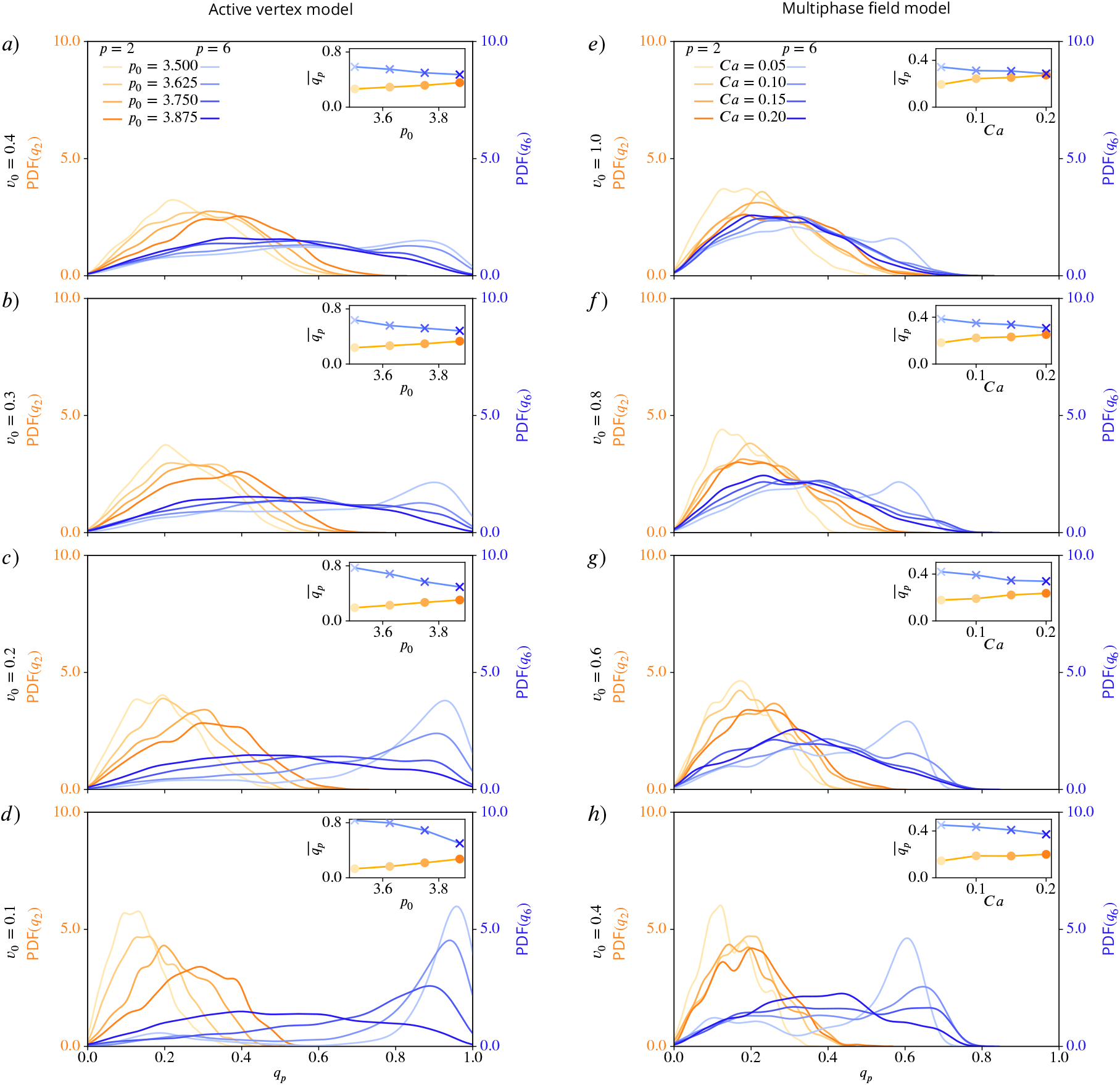
Nematic (*p* = 2) and hexatic (*p* = 6) order depend on deformability of the cells. Probability distribution functions (PDFs) for *q*_2_ (shades of orange) and *q*_6_ (shades of blue), using kde-plots, for varying deformability *p*_0_ or *Ca* and fixed activity *v*_0_. Inlets show mean values of *q*_2_ and *q*_6_ as function of deformability. a) - d) active vertex model, e) - h) multiphase field model for decreasing activity.

**Figure 8—figure supplement 2.**
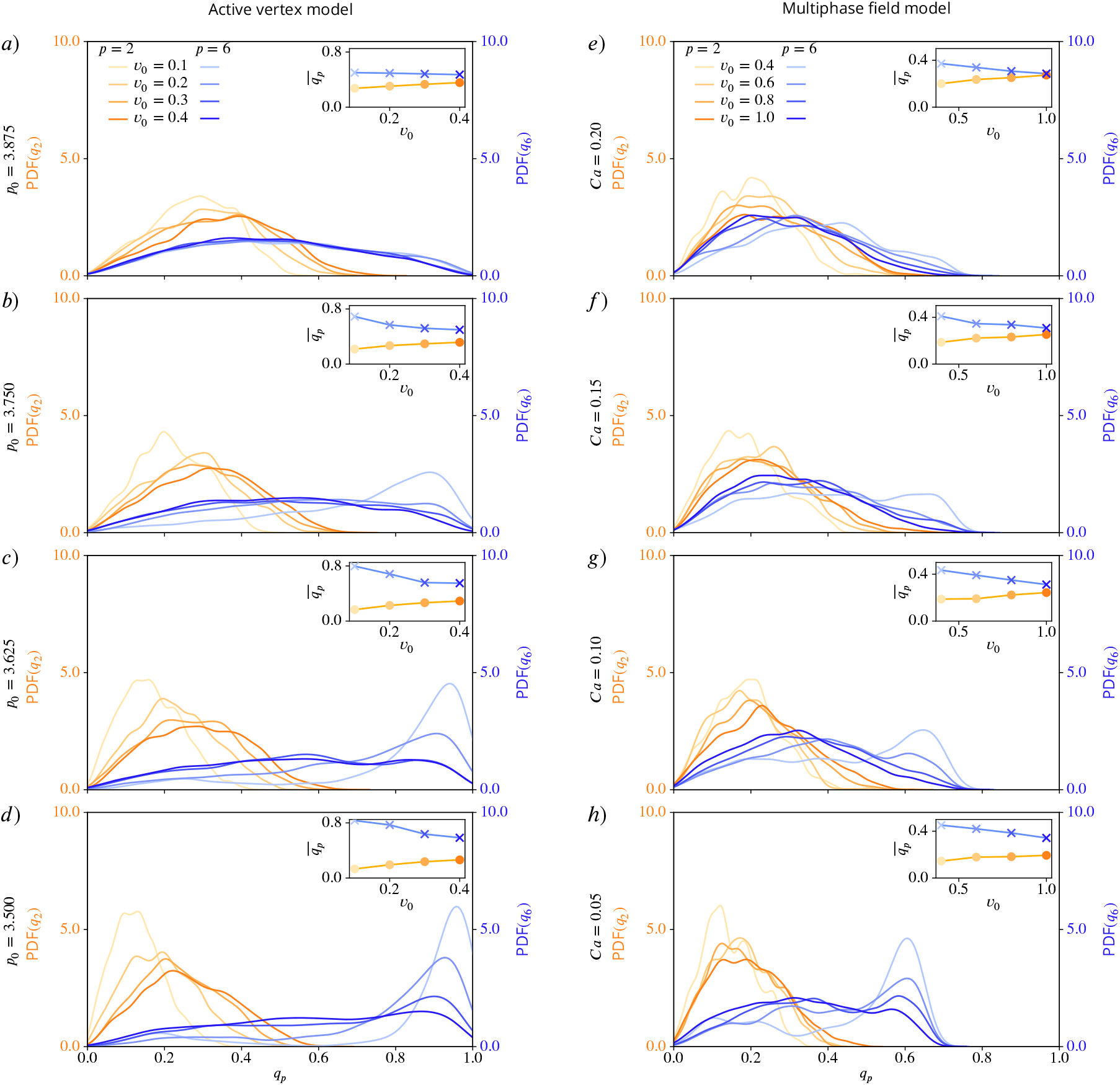
Nematic (*p* = 2) and hexatic (*p* = 6) order depend on activity of the cells. Probability distribution functions (PDFs) for *q*_2_ (shades of orange) and *q*_6_ (shades of blue), using kde-plots, for varying activity *v*_0_ and fixed deformability *p*_0_ or *Ca*. Inlets show mean values of *q*_2_ and *q*_6_ as function of activity. a) - d) active vertex model and e) - h) multiphase field model for decreasing deformability.

**Figure 9—figure supplement 1.**
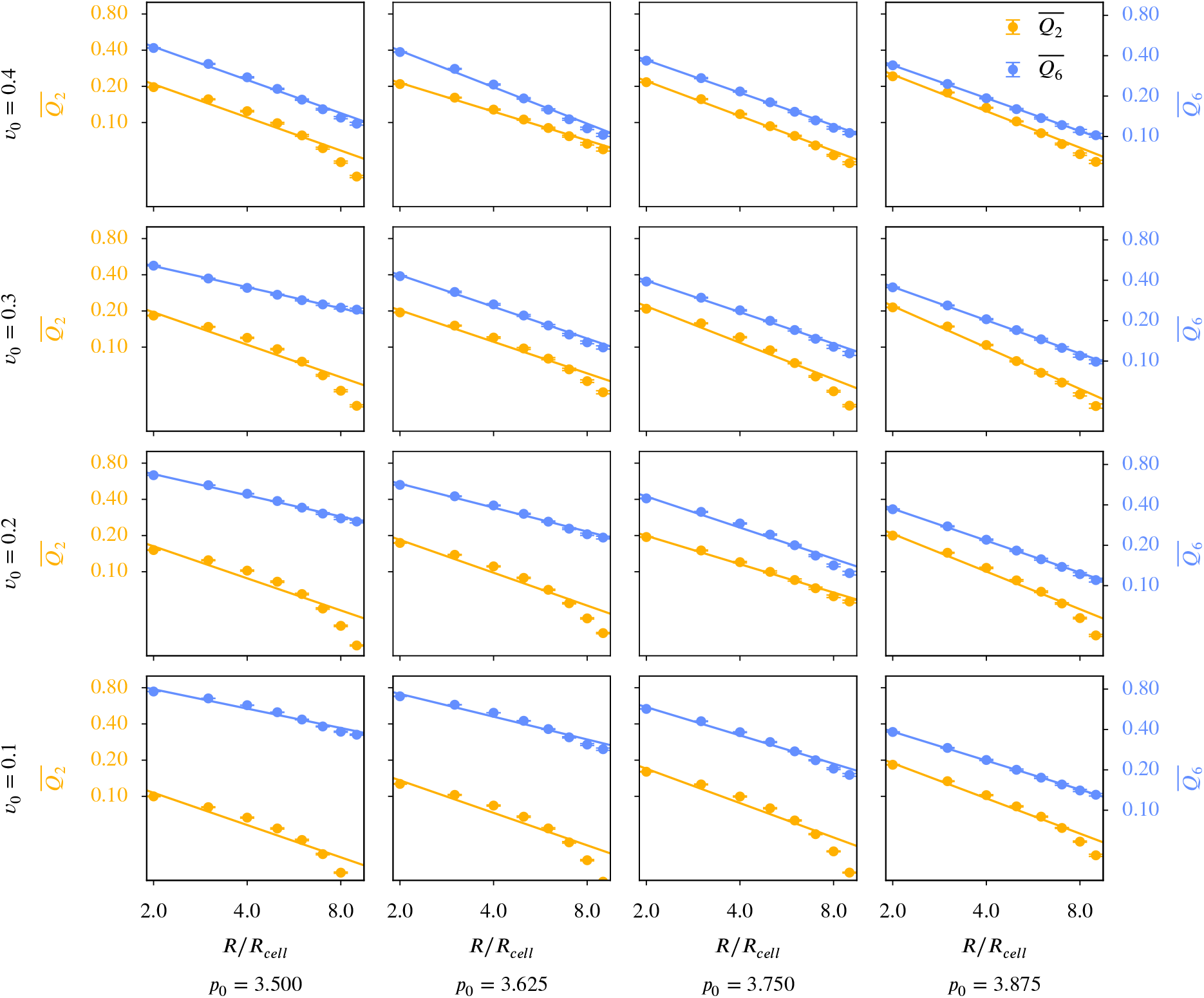
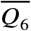 versus 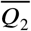 for different coarse graining radii in the active vertex model. *Q*_*p*_ was calculated according to ***Equation 11***, the averaging of this and the choice of *R*_𝒞_ follow the description in Coarse-grained quantities. The maximal coarse-graining radius corresponds to half the domain width. Each panel corresponds to specific model parameters; *p*_0_ and *v*_0_, representing deformability and activity. A logarithmic scaling was used for both axis. Error bars are obtained as s.e.m..

**Figure 9—figure supplement 2.**
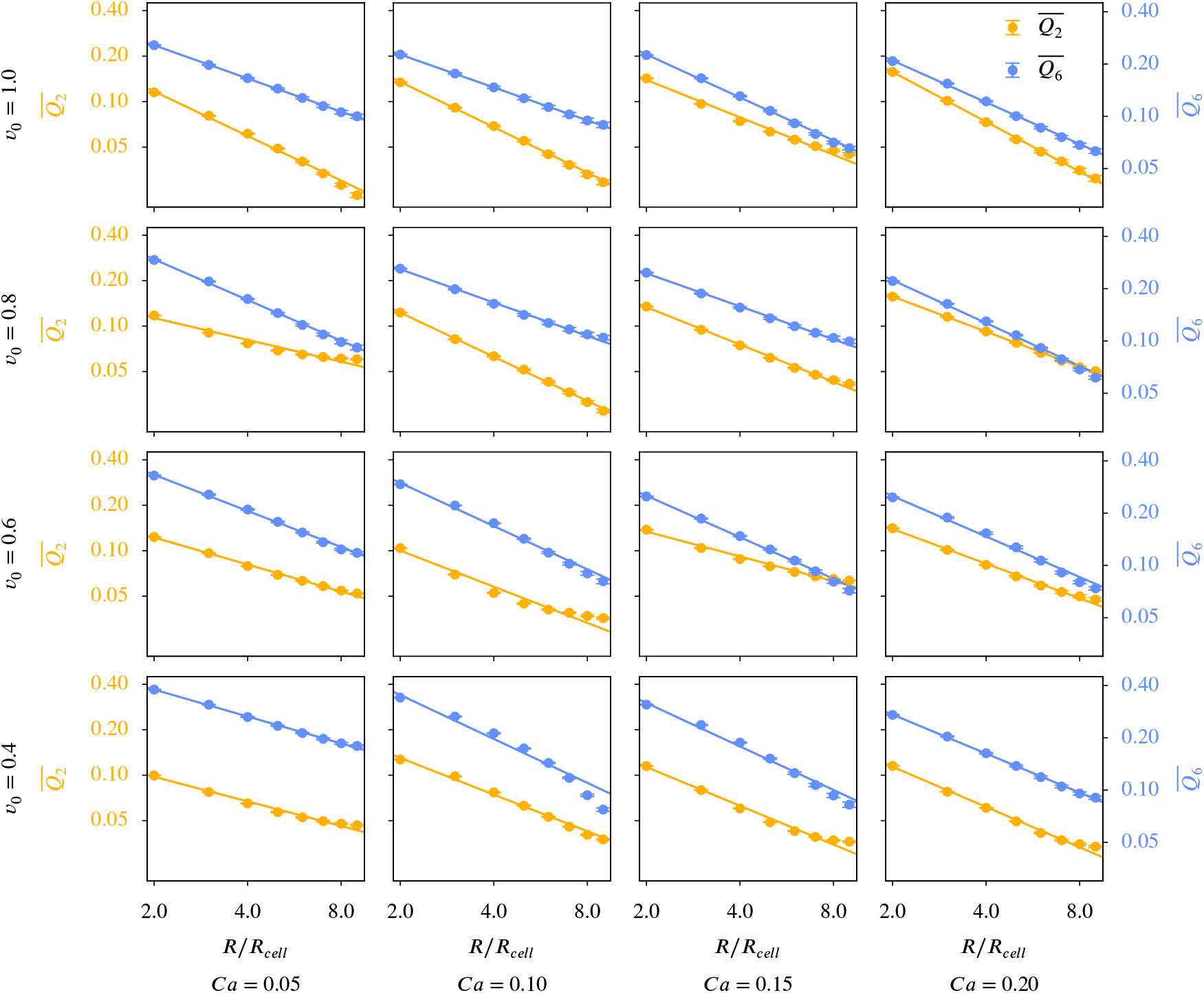
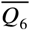 versus 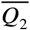 for different coarse graining radii in the multiphase field model. *Q*_*p*_ was calculated according to ***Equation 11***, the averaging of this and the choice of *R*_𝒞_ follow the description in Coarse-grained quantities. The maximal coarse-graining radius corresponds to half the domain width. Each panel corresponds to specific model parameters; *Ca* and *v*_0_, representing deformability and activity. A logarithmic scaling was used for both axis. Error bars are obtained as s.e.m..

**Figure 10—figure supplement 1.**
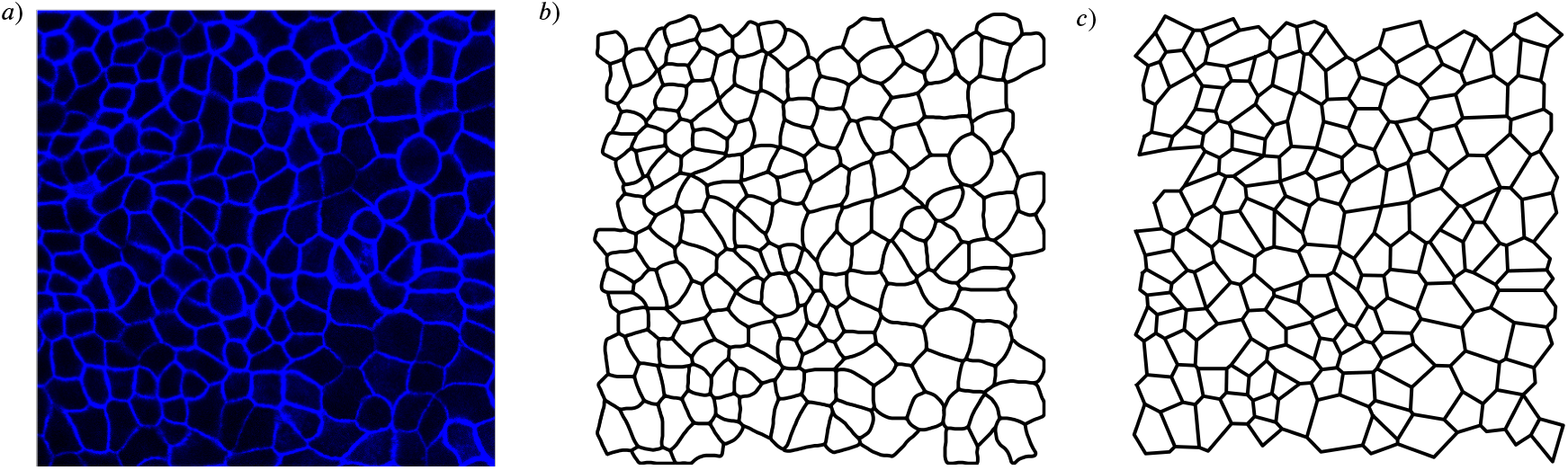
Cell shapes in the experiments from ***Armengol-Collado et al. (2023***) *a*) and the cell shapes used for the shape quantification, b) detailed cell outline obtained from the microscopy image, c) polygonal approximation of the cell shape. Shown for frame number 1.

**Figure 10—figure supplement 2.**
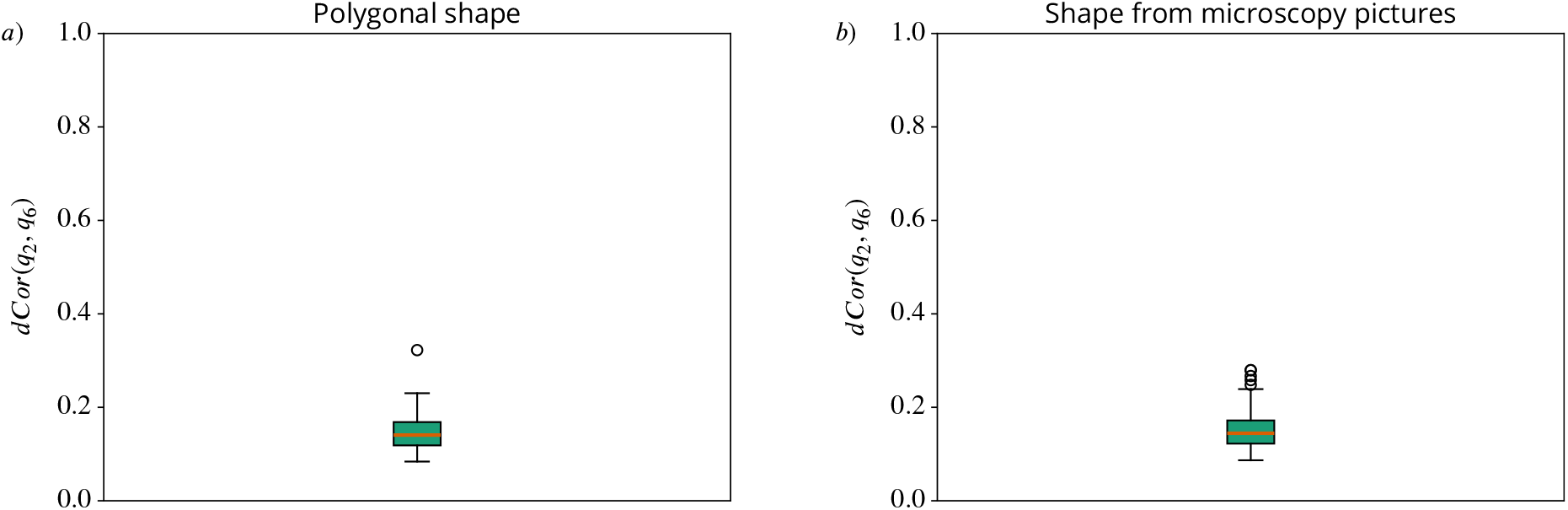
Distance correlation *dCor*(*q*_2_, *q*6) between *q*_2_ and *q*_6_. On the left side *a*) we use the polygonal approximation of the cell shape, on the right side *b*) we use the detailed cell outline obtained from the microscopy pictures. We compute one value for every frame and present the resulting distribution with a box plot. In the box plots the orange line corresponds to the median of the data and the box ranges from the first to the third quartile of the data. The whiskers go from the lowest data point greater than (value of the first quartile) − 1.5 × (Interquartile range) to the highest data point below (value of the third quartile) + 1.5 × (Interquartile range). Outliers are shown with circles.

**Figure 10—figure supplement 3.**
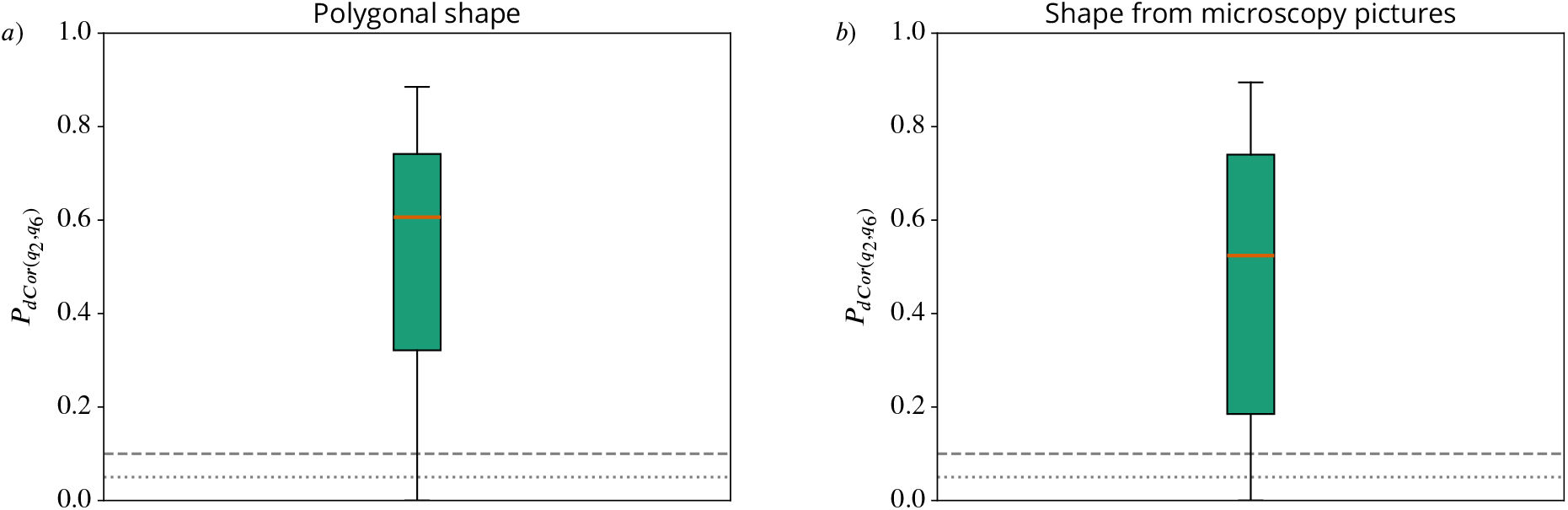
P-values of the distance correlation 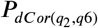 between *q*_2_ and *q*_6_. On the left side *a*) we use the polygonal approximation of the cell shape, on the right side *b*) we use the detailed cell outline obtained from the microscopy pictures. We compute one value for every frame and present the resulting distribution with a box plot. In the box plots the orange line corresponds to the median of the data and the box ranges from the first to the third quartile of the data. The whiskers go from the lowest data point greater than (value of the first quartile) − 1.5 × (Interquartile range) to the highest data point below (value of the third quartile) + 1.5 × (Interquartile range). Outliers are shown with circles. 0.05 and 0.1 are marked with a grey dotted/grey dashed line to guide the eye.

**Appendix 3—figure 24—figure supplement 1.**
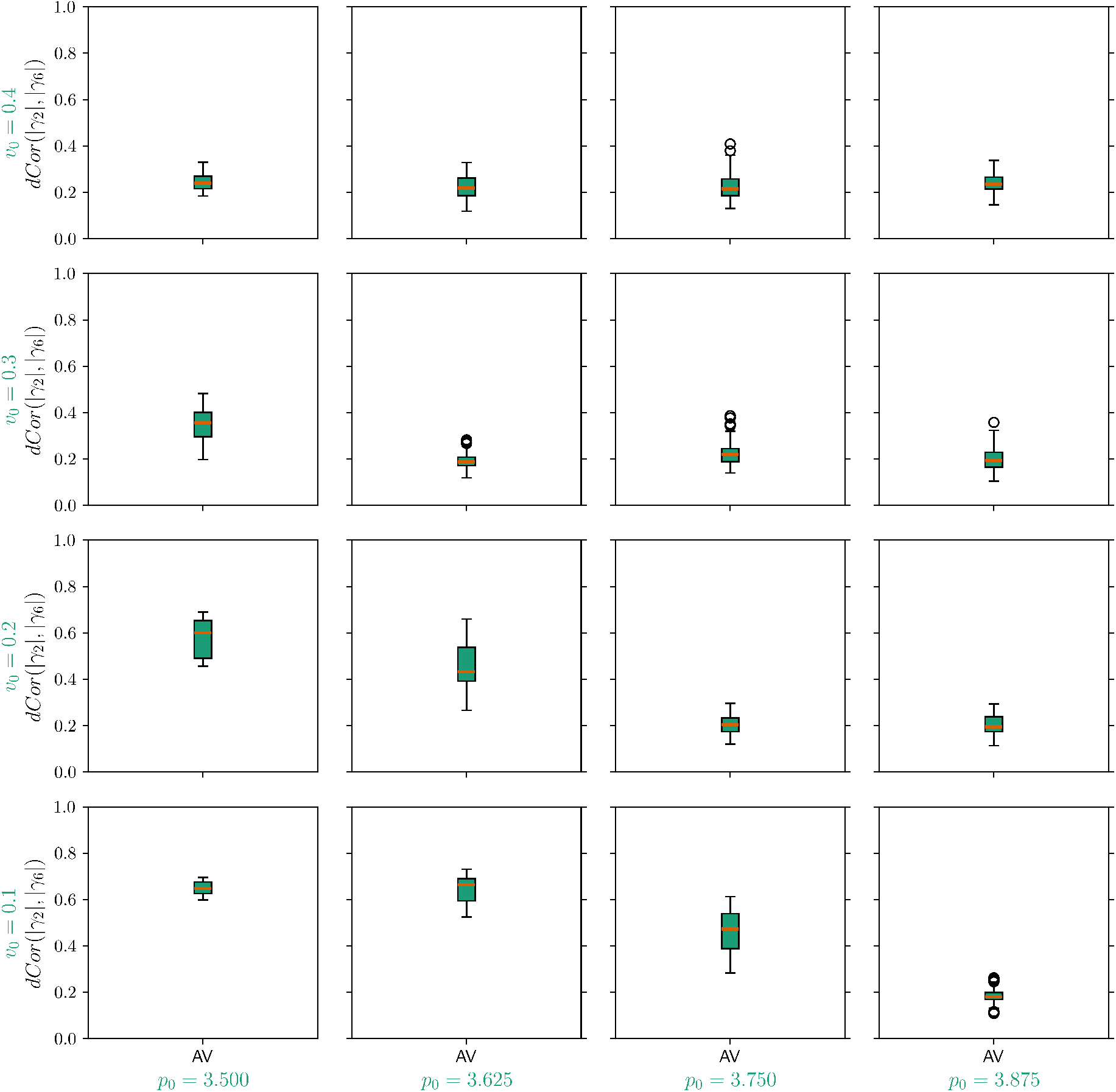
Distance correlation *dCor*(|*γ*_2_|, | *γ*_6_|) for the simulation data of the active vertex model. Each panel corresponds to specific model parameters *p*_0_ and *v*_0_, representing deformability and activity. We compute one value for every timestep and present the resulting distribution with a box plot. In the box plots the orange line corresponds to the median of the data and the box ranges from the first to the third quartile of the data. The whiskers go from the lowest data point greater than (value of the first quartile) − 1.5 × (Interquartile range) to the highest data point below (value of the third quartile) + 1.5 × (Interquartile range). Outliers are shown with circles.

**Appendix 3—figure 24—figure supplement 2.**
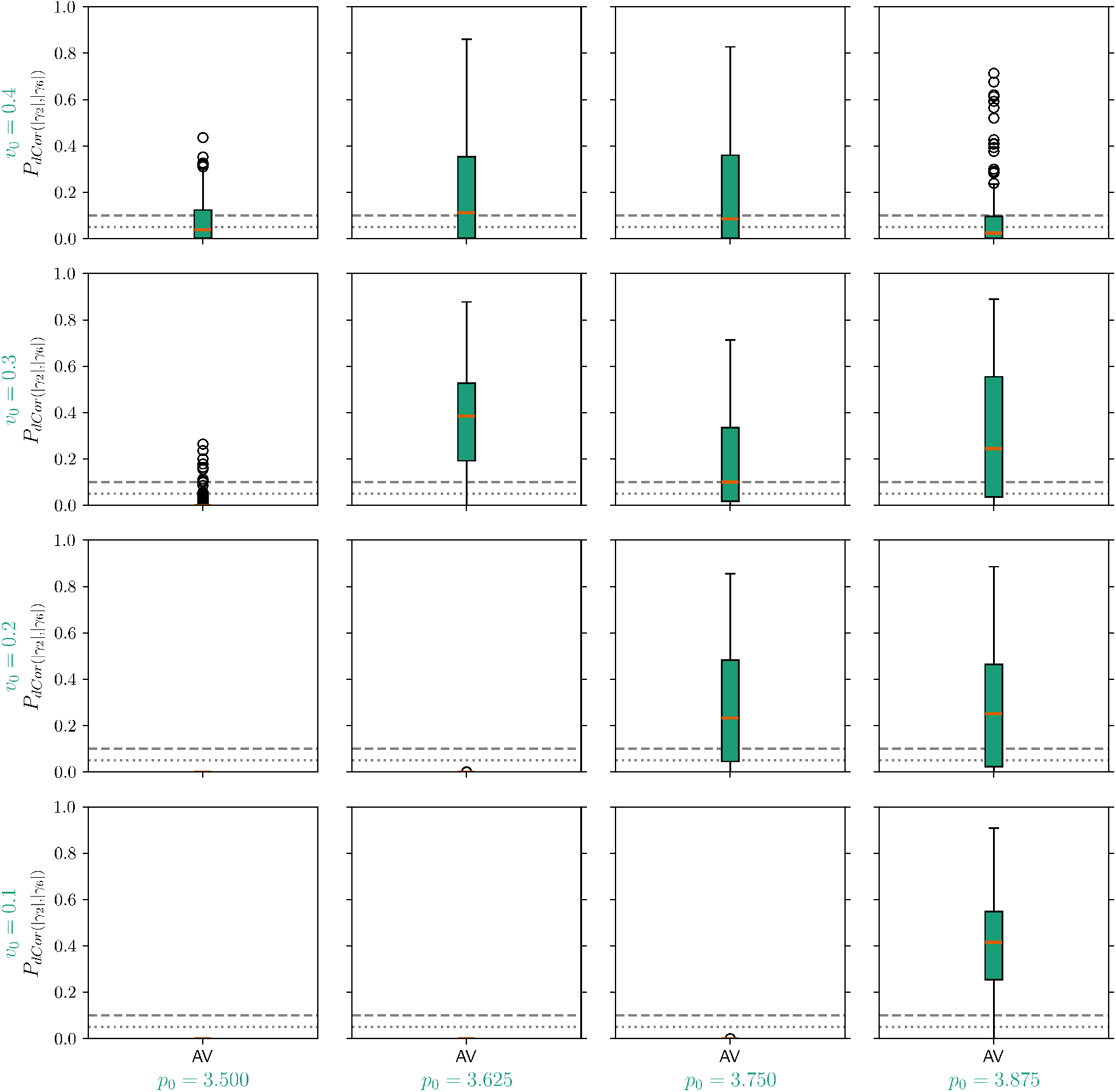
P-values of the distance correlation 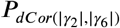 for the simulation data of the active vertex model. Each panel corresponds to specific model parameters *p*_0_ and *v*_0_, representing deformability and activity. We compute one value for every timestep and present the resulting distribution with a box plot. In the box plots the orange line corresponds to the median of the data and the box ranges from the first to the third quartile of the data. The whiskers go from the lowest data point greater than (value of the first quartile) − 1.5 × (Interquartile range) to the highest data point below (value of the third quartile) + 1.5 × (Interquartile range). Outliers are shown with circles. 0.05 and 0.1 are marked with a grey dotted/grey dashed line to guide the eye.

**Appendix 3—figure 25—figure supplement 1.**
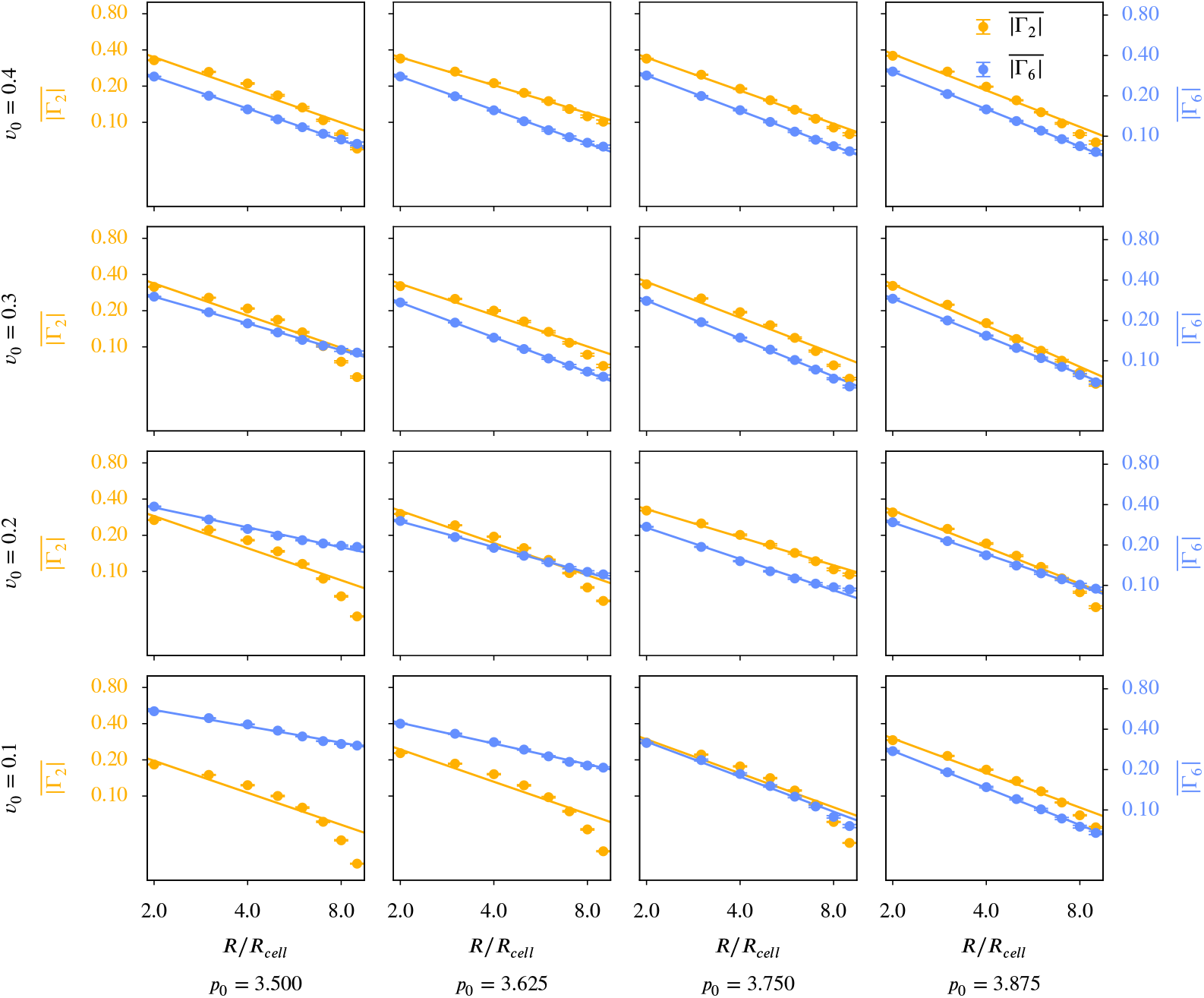
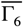 versus 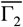 for different coarse graining radii in the active vertex model. Γ_*p*_ was calculated according to ***Equation 12***, the averaging of this and the choice of *R*_𝒞_ follow the description in Coarse-grained quantities. The maximal coarse-graining radius corresponds to half the domain width. Each panel corresponds to specific model parameters; *p*_0_ and *v*_0_, representing deformability and activity. A logarithmic scaling was used for both axis. Error bars are obtained as s.e.m..

**Appendix 3—figure 26—figure supplement 1.**
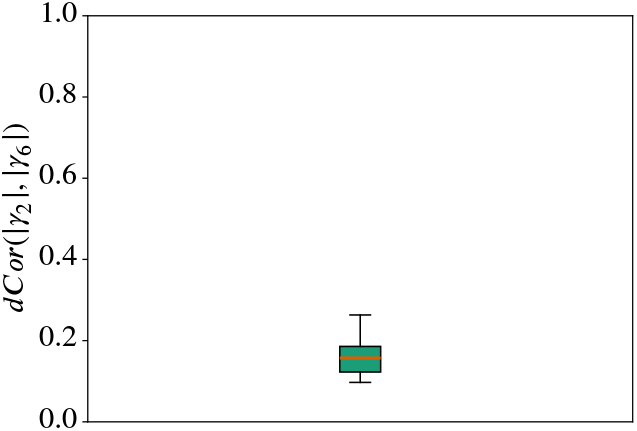
Distance correlation *dCor*(|*γ*_2_|, |*γ*_6_|) for all cells from the experimental data in ***Armengol-Collado et al. (2023***). We use the polygonal approximation of the cell shape as *γ*_*p*_ can only work with polygons. We compute one value for every frame and present the resulting distribution with a box plot. In the box plots the orange line corresponds to the median of the data and the box ranges from the first to the third quartile of the data. The whiskers go from the lowest data point greater than (value of the first quartile) − 1.5 × (Interquartile range) to the highest data point below (value of the third quartile) + 1.5 × (Interquartile range). Outliers are shown with circles.

**Appendix 3—figure 26—figure supplement 2.**
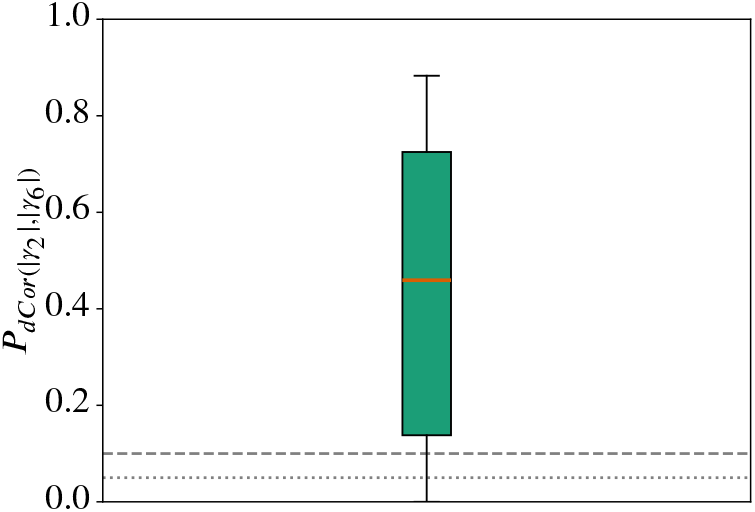
P-value of the distance correlation 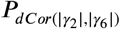 for all cells from the experimental data in ***Armengol-Collado et al. (2023***). We use the polygonal approximation of the cell shape as *γ*_*p*_ can only work with polygons. We compute one value for every frame and present the resulting distribution with a box plot. In the box plots the orange line corresponds to the median of the data and the box ranges from the first to the third quartile of the data. The whiskers go from the lowest data point greater than (value of the first quartile) − 1.5 × (Interquartile range) to the highest data point below (value of the third quartile) + 1.5 × (Interquartile range). Outliers are shown with circles.

## References

Alert R, Trepat X. Physical models of collective cell migration. Annual Review of Condensed Matter Physics. 2020; 11:77–101.

Alesker S. Description of continuous isometry covariant valuations on convex sets. Geometriae Dedicata. 1999; 74:241–248.

Alt S, Ganguly P, Salbreux G. Vertex models: from cell mechanics to tissue morphogenesis. Philosophical Transactions of the Royal Society B: Biological Sciences. 2017; 372(1720):20150520.

Armengol-Collado JM, Carenza LN, Eckert J, Krommydas D, Giomi L. Epithelia are multiscale active liquid crys-tals. Nature Physics. 2023; 19:1773–1779.

Armengol-Collado JM, Carenza LN, Giomi L. Hydrodynamics and multiscale order in confluent epithelia. eLife. 2024; 13:e86400.

Balasubramaniam L, Doostmohammadi A, Saw TB, Narayana GHNS, Mueller R, Dang T, Thomas M, Gupta S, Sonam S, Yap AS, Toyama Y, Mège RM, Yeomans JM, Ladoux B. Investigating the nature of active forces in tissues reveals how contractile cells can form extensile monolayers. Nature Materials. 2021; 20(8):1156– 1166.

Beisbart C, Barbosa MS, Wagner H, Costa LdF. Extended morphometric analysis of neuronal cells with minkowski valuations. The European Physical Journal B. 2006; 52:531–546.

Beisbart C, Dahlke R, Mecke K, Wagner H. Vector- and tensor-valued descriptors for spatial patterns. In: Mecke K, Stoyan D, editors. Morphology of Condensed Matter: Physics and Geometry of Spatially Complex Systems Berlin, Heidelberg: Springer Berlin Heidelberg; 2002. p. 238–260.

Bernard EP, Krauth W. Two-step melting in two dimensions: first-order liquid-hexatic transition. Physical Review Letters. 2011; 107:155704.

Bi D, Yang X, Marchetti MC, Manning ML. Motility-driven glass and jamming transitions in biological tissues. Physical Review X. 2016; 6:021011.

Bi D, Lopez J, Schwarz JM, Manning ML. A density-independent rigidity transition in biological tissues. Nature Physics. 2015; 11(12):1074–1079.

Bladon P, Frenkel D. Dislocation unbinding in dense two-dimensional crystals. Physical Review Letters. 1995; 74:2519–2522.

Bowick MJ, Manyuhina OV, Serafin F. Shapes and singularities in triatic liquid-crystal vesicles. Europhysics Letters. 2017; 117(2):26001.

Cislo DJ, Yang F, Qin H, Pavlopoulos A, Bowick MJ, Streichan SJ. Active cell divisions generate fourfold orienta-tionally ordered phase in living tissue. Nature Physics. 2023; 19(8):1201–1210.

Das A, Sastry S, Bi D. Controlled neighbor exchanges drive glassy behavior, intermittency, and cell streaming in epithelial tissues. Physical Review X. 2021; 11(4):041037.

Duclos G, Erlenkämper C, Joanny JF, Silberzan P. Topological defects in confined populations of spindle-shaped cells. Nature Physics. 2017; 13(1):58–62.

Durand M, Heu J. Thermally driven order-disorder transition in two-dimensional soft cellular systems. Physical Review Letters. 2019; 123(18):188001.

Eckert J, Ladoux B, Mège RM, Giomi L, Schmidt T. Hexanematic crossover in epithelial monolayers depends on cell adhesion and cell density. Nature Communications. 2023; 14(1):5762.

Farhadifar R, Röper JC, Aigouy B, Eaton S, Jülicher F. The influence of cell mechanics, cell-cell interactions, and proliferation on epithelial packing. Current biology. 2007; 17(24):2095–2104.

Fletcher AG, Osterfield M, Baker RE, Shvartsman SY. Vertex models of epithelial morphogenesis. Biophysical Journal. 2014; 106(11):2291–2304.

Gasser U, Eisenmann C, Maret G, Keim P. Melting of crystals in two dimensions. ChemPhysChem. 2010; 11(5):963–970.

de Gennes PG, Prost J. The physics of liquid crystals. Oxford: Second Edition, Clarendon Press; 1993.

Giomi L, Toner J, Sarkar N. Hydrodynamic theory of p-atic liquid crystals. Physical Review E. 2022; 106(2):024701.

Giomi L, Toner J, Sarkar N. Long-ranged order and flow alignment in sheared p-atic liquid crystals. Physical Review Letters. 2022; 129(6):067801.

Graner F, Dollet B, Raufaste C, Marmottant P. Discrete rearranging disordered patterns, part I: robust statistical tools in two or three dimensions. The European physical journal E, Soft matter. 2008; 25(4):349–369.

Hadwiger H. Vorlesungen über Inhalt, Oberfläche und Isoperimetrie. Springer Berlin Heidelberg; 1975.

Hakim V, Silberzan P. Collective cell migration: a physics perspective. Reports on Progress in Physics. 2017; 80:076601.

Halperin BI, Nelson DR. Theory of two-dimensional melting. Physical Review Letters. 1978; 41:121–124.

Happel L, Voigt A. Coordinated motion of epithelial layers on curved surfaces. Physical Review Letters. 2024; 132:078401.

Honda H. Geometrical models for cells in tissues. International review of cytology. 1983; 81:191–248.

Honda H, Nagai T. Mathematical models of cell-based morphogenesis. Springer; 2022.

Hug D, Schneider R, Schuster R. The space of isometry covariant tensor valuations. St Petersburg Mathematical Journal. 2007 02; 19:137–158.

Hug D, Schneider R, Schuster R. Integral geometry of tensor valuations. Advances in Applied Mathematics. 2008; 41(4):482–509.

Jain HP, Hu RDJG, Angheluta L. Emergent directional migration due to interplay between cell shape and T1 transitions. Under review. 2024;.

Jain HP, Voigt A, Angheluta L. Robust statistical properties of T1 transitions in a multi-phase field model of cell monolayers. Scientific Reports. 2023; 13(1):10096.

Jain HP, Voigt A, Angheluta L. From cell intercalation to flow, the importance of T1 transitions. Physical Review Research. 2024; 6(3):033176.

Kapfer S. Morphometry and Physics of Particulate and Porous Media. PhD thesis, FAU Erlangen-Nürnberg; 2012.

Kapfer SC, Mickel W, Mecke K, Schröder-Turk GE. Jammed spheres: Minkowski tensors reveal onset of local crystallinity. Physical Review E. 2012; 85:030301.

Kawaguchi K, Kageyama R, Sano M. Topological defects control collective dynamics in neural progenitor cell cultures. Nature. 2017; 545(7654):327–331.

Killeen A, Bertrand T, Lee CF. Polar fluctuations lead to extensile nematic behavior in confluent tissues. Physical Review Letters. 2022; 128(7):078001.

Klain DA. The minkowski problem for polytopes. Advances in Mathematics. 2004; 185(2):270–288.

Klatt MA, Hörmann M, Mecke K. Characterization of anisotropic gaussian random fields by minkowski tensors. Journal of Statistical Mechanics: Theory and Experiment. 2022; 2022(4):043301.

Koride S, Loza AJ, Sun SX. Epithelial vertex models with active biochemical regulation of contractility can explain organized collective cell motility. APL bioengineering. 2018; 2(3).

Krommydas D, Carenza LN, Giomi L. Hydrodynamic enhancement of p-atic defect dynamics. Physical Review Letters. 2023; 130(9):098101.

Li X, Das A, Bi D. Mechanical heterogeneity in tissues promotes rigidity and controls cellular invasion. Physical Review Letters. 2019; 123:058101.

Li YW, Ciamarra MP. Role of cell deformability in the two-dimensional melting of biological tissues. Physical Review Materials. 2018; 2(4):045602.

Loewe B, Chiang M, Marenduzzo D, Marchetti MC. Solid-liquid transition of deformable and overlapping active particles. Physical Review Letters. 2020; 125:038003.

Maroudas-Sacks Y, Garion L, Shani-Zerbib L, Livshits A, Braun E, Keren K. Topological defects in the nematic order of actin fibres as organization centres of hydra morphogenesis. Nature Physics. 2021; 17(2):251–259.

McMullen P. Isometry covariant valuations on convex bodies. In: Second international conference in stochastic geometry, convex bodies and empirical measures, Agrigento, Italy, September 9–14, 1996 Palermo: Circolo Matematico di Palermo; 1997.p. 259–271.

Mecke KR. Additivity, convexity, and beyond: Applications of minkowski functionals in statistical physics. In: Mecke KR, Stoyan D, editors. Statistical Physics and Spatial Statistics Berlin, Heidelberg: Springer Berlin Heidelberg; 2000. p. 111–184.

Melo S, Guerrero P, Moreira Soares M, Bordin JR, Carneiro F, Carneiro P, Dias MB, Carvalho J, Figueiredo J, Seruca R, Travasso RDM. The ECM and tissue architecture are major determinants of early invasion mediated by E-cadherin dysfunction. Communications Biology. 2023; 6(1):1132.

Merkel M, Etournay R, Popović M, Salbreux G, Eaton S, Jülicher F. Triangles bridge the scales: Quantifying cellular contributions to tissue deformation. Physical Review E. 2017; 95(3):032401.

Mickel W, Kapfer SC, Schröder-Turk GE, Mecke K. Shortcomings of the bond orientational order parameters for the analysis of disordered particulate matter. The Journal of Chemical Physics. 2013; 138(4):044501.

Minkowski H. Allgemeine Lehrsätze über die convexen Polyeder. Nachrichten von der Gesellschaft der Wissenschaften zu Göttingen, Mathematisch-Physikalische Klasse. 1897; 1897:198–220.

Monfared S, Ravichandran G, Andrade J, Doostmohammadi A. Mechanical basis and topological routes to cell elimination. eLife. 2023; 12:e82435.

Moure A, Gomez H. Phase-field modeling of individual and collective cell migration. Archives of Computational Methods in Engineering. 2021; 28:311–344.

Mueller R, Yeomans JM, Doostmohammadi A. Emergence of active nematic behavior in monolayers of isotropic cells. Physical Review Letters. 2019; 122:048004.

Murray CA, Van Winkle DH. Experimental observation of two-stage melting in a classical two-dimensional screened coulomb system. Physical Review Letters. 1987 Mar; 58:1200–1203.

Nelson DR, Halperin BI. Dislocation-mediated melting in two dimensions. Physical Review B. 1979 Mar; 19:2457–2484.

Onsager L. The effects of shape on the interaction of colloidal particales. Annals of the New York Academy of Sciences. 1949; 51(4):627–659.

Pasupalak A, Yan-Wei L, Ni R, Ciamarra MP. Hexatic phase in a model of active biological tissues. Soft matter. 2020; 16(16):3914–3920.

Praetorius S, Voigt A. Collective cell behaviour – a cell-based parallelisation approach for a phase field active polar gel model. No. 49 in NIC Series, Jülich: Forschungszentrum Jülich GmbH, Zentralbibliothek; 2018.

Ravichandran Y, Vogg M, Kruse K, Pearce DJ, Roux A. Topology changes of the regenerating hydra define actin nematic defects as mechanical organizers of morphogenesis. bioRxiv. 2024; p. 2024–04.

Salvalaglio M, Voigt A, Wise SM. Doubly degenerate diffuse interface models of surface diffusion. Mathematical Methods in the Applied Sciences. 2021; 44:5385–5405.

Saw TB, Doostmohammadi A, Nier V, Kocgozlu L, Thampi S, Toyama Y, Marcq P, Lim CT, Yeomans JM, Ladoux B. Topological defects in epithelia govern cell death and extrusion. Nature. 2017; 544(7649):212–216.

Saye RI, Sethian JA. Analysis and applications of the voronoi implicit interface method. Journal of Computational Physics. 2012; 231(18):6051–6085.

Saye RI, Sethian JA. The voronoi implicit interface method for computing multiphase physics. Proceedings of the National Academy of Science. 2011; 108(49):19498–19503.

Schaller F, Kapfer S, Klatt M, Hug D, Last G, Mecke K, Schröder-Turk G, morphometry.org; 2024. https://morphometry.org.

Schneider R. Convex bodies: The Brunn–Minkowski theory. Encyclopedia of Mathematics and its Applications, Cambridge University Press; 2013.

Schröder-Turk GE, Mickel W, Kapfer SC, Schaller FM, Breidenbach B, Hug D, Mecke K. Minkowski tensors of anisotropic spatial structure. New Journal of Physics. 2013; 15(8):083028.

Schröder-Turk GE, Mickel W, Schröter M, Delaney GW, Saadatfar M, Senden TJ, Mecke K, Aste T. Disordered spherical bead packs are anisotropic. Europhysics Letters. 2010; 90(3):34001.

Schröder-Turk GE, Kapfer S, Breidenbach B, Beisbart C, Mecke K. Tensorial minkowski functionals and anisotropy measures for planar patterns. Journal of Microscopy. 2010; 238(1):57–74.

Sknepnek R, Djafer-Cherif I, Chuai M, Weijer C, Henkes S. Generating active T1 transitions through mechanochemical feedback. eLife. 2023; 12:e79862.

Stringer C, Wang T, Michaelos M, Pachitariu M. Cellpose: a generalist algorithm for cellular segmentation. Nature Methods. 2021; 18:100–106.

Székely GJ, Rizzo ML, Bakirov NK. Measuring and testing dependence by correlation of distances. The Annals of Statistics. 2007; 35(6):2769–2794.

Tong S, Sknepnek R, Košmrlj A. Linear viscoelastic response of the vertex model with internal and external dissipation: Normal modes analysis. Physical Review Research. 2023; 5(1):013143.

Vaxman A, Campen M, Diamanti O, Panozzo D, Bommes D, Hildebrandt K, Ben-Chen M. Directional field synthesis, design, and processing. Computer Graphics Forum. 2016; 35(2):545–572.

Vey S, Voigt A. AMDiS: Adaptive multidimensional simulations. Computing and Visualization in Science. 2007; 10(1):57–67.

Virga EG. Octupolar order in two dimensions. The European Physical Journal E. 2015; 38(6).

van der Walt S, Schönberger JL, Nunez-Iglesias J, Boulogne F, Warner JD, Yager N, Gouillart E, Yu T, the scikitimage contributors. scikit-image: image processing in Python. PeerJ. 2014; 2:e453. doi: 10.7717/peerj.453.

Wang PY, Mason TG. A Brownian quasi-crystal of pre-assembled colloidal penrose tiles. Nature. 2018; 561(7721):94–99.

Wenzel D, Voigt A. Multiphase field models for collective cell migration. Physical Review E. 2021; 184:054410.

Witkowski T, Ling S, Praetorius S, Voigt A. Software concepts and numerical algorithms for a scalable adaptive parallel finite element method. Advances in Computational Mathematics. 2015; 41(6):1145–1177.

Wojciechowski KW, Frenkel D. Tetratic phase in the planar hard square system? Computational Methods in Science and Technology. 2004; 10:235–255.

Yu T, Mason TG. Heptatic liquid quasi-crystals by colloidal lithographic pre-assembly. Journal of Colloid and Interface Science. 2023;.

Zahn K, Lenke R, Maret G. Two-stage melting of paramagnetic colloidal crystals in two dimensions. Physical Review Letters. 1999; 82:2721–2724.

